# Sensitivity of genome-wide tests for mitonuclear genetic incompatibilities

**DOI:** 10.1101/2025.06.30.662443

**Authors:** Shady A. Kuster, Molly Schumer, Justin C. Havird, Daniel B. Sloan

## Abstract

Mismatches between interacting mitochondrial and nuclear gene products in hybrids have been proposed to disproportionately contribute to early species boundaries. Under this model, genetic incompatibilities emerge when mitochondrial haplotypes are in a cellular context without their coevolved nuclear-encoded mitochondrial (n-mt) proteins. Some case studies have shown that such disruptions in mitonuclear coevolution can contribute to reproductive isolation, but whether mitonuclear incompatibilities generate selection that impacts multiple n-mt loci and/or causes broad, genome-wide contributions to speciation is unclear. Here, we leverage a system with several hybridizing species pairs (*Xiphophorus* fishes) that have known mitonuclear incompatibilities of large effect. We divided nuclear-encoded genes into three classes based on level of interaction with mitochondrial gene products. We found only inconsistent statistical support for a difference between these classes in the degree of positive covariation in mitonuclear ancestry. We discuss evidence that these analyses are sensitive to the amount of non-synonymous divergence between parent species in interacting n-mt genes or the age of the hybridization event. Overall, our results imply that genome-wide scans focused on enrichment of broad functional gene classes may often be insufficient for detecting a history of mitonuclear coevolution, even when strong selection is acting on mitonuclear incompatibilities at multiple loci.

## INTRODUCTION

Since its endosymbiotic origin, the mitochondrion has evolved into an integral component of the eukaryotic cell, functioning in such processes as oxidative phosphorylation (OXPHOS), Ca^2+^ acquisition and signaling, and biosynthesis of heme and iron-sulfur clusters. Bilaterian metazoan mitochondrial genomes (mitogenomes) have been reduced to encode only 13 core protein subunits of the electron transport chain, 2 rRNAs, and 22 tRNAs, making mitochondria dependent on the import of nuclear-encoded, but mitochondrially targeted, (n-mt) proteins for all functions [1–4]. Due to the importance of mitochondrial function to fitness, maintaining these intimate interactions among coevolved partners may be essential.

Mitogenomes evolve at a faster rate than nuclear genomes in most bilaterian metazoan systems, with an average rate of 19× faster in vertebrates [5–8]. These elevated substitution rates may result from a higher underlying mutation rate in combination with inefficient purging of deleterious mutations due to smaller effective population size, lack of recombination, and uniparental inheritance [9–12], although explanations based on reduced efficacy of selection in mitogenomes have been recently challenged [13–15]. Regardless of the cause, the consequences of high mitochondrial DNA (mtDNA) substitution rates are often deleterious [6]. As such, substitutions in mitochondrially encoded proteins may create selection for compensatory changes in interacting n-mt proteins [16–18], and signatures of coevolution have indeed been found between the mitochondrial and nuclear genomes [19–22]. Other forms of mitonuclear coevolution (i.e., reciprocal selection for compatible genotypes) are also possible [19,23–25]. Regardless of the exact process, the outcome is a set of interacting proteins that are functionally dependent on sufficient “matching” of lineage-specific substitutions in mitochondrial and n-mt proteins [23].

As a result, disrupting the association between a mitochondrial haplotype and its coevolved nuclear counterparts (e.g., via secondary contact following speciation) is predicted to uncover genetic incompatibilities that made early contributions to reproductive isolation [5,26]. Given the rate at which mitonuclear genes coevolve and the importance of mitochondrial function to cellular metabolism, mitonuclear incompatibilities are proposed to have a disproportionate effect on the accumulation of reproductive barriers during lineage divergence [23,27–30]. These incompatibilities could potentially be driven by only a few genes with major effects on fitness or alternatively by many genes of small effect. Although studies have found evidence of mitonuclear incompatibilities (e.g., [31,32]) in some systems, it is still unclear how far-reaching these effects are and how many genes are involved. For example, the immediate effects of selection on mitonuclear incompatibilities could be restricted to a subset of n-mt proteins involved in direct physical interactions with mitochondrial gene products. On the other hand, because mitochondria are hubs of various biological processes, selection may be more widespread and affect nuclear genes that have no direct connection to mitochondrial function. We hypothesize that, relative to other nuclear genes, n-mt genes are more strongly affected by selection acting to maintain associations between matched lineage-specific mitonuclear genotypes. Therefore, examining signatures of mtDNA “match” in n-mt genes vs. nuclear-encoded genes that lack mitochondrial interactions should reveal a history of mitonuclear coevolution across the genome at species boundaries. While partitioning genomic data based on n-mt gene function has been used to investigate mitonuclear incompatibilities in many systems (Table S1), the effectiveness of this approach has not been tested in a system with previously known mitonuclear incompatibilities.

Species of the swordtail fish genus *Xiphophorus* have recently begun hybridizing in multiple river systems in Mexico, providing a model of repeated hybridization and speciation dynamics [33–42]. Recent work on hybrid zones between sister species *X. birchmanni* and *X. malinche* has identified severe mitonuclear incompatibilities involving the *ndufs5* and *ndufa13* n-mt genes [43]. Mitonuclear ancestry mismatches at these loci have severe fitness effects, including death during embryonic development. Subsequent investigations have identified that at least nine nuclear genomic regions (which contain a total of 10 known n-mt genes across seven loci) are involved in mitonuclear incompatibilities in these hybrids [44]. Some of these incompatibilities, including the *ndufs5* and *ndufa13* n-mt genes, were also found in *X. birchmanni* and *X. cortezi* hybrids [42]. We therefore leveraged this system to investigate evidence for mitonuclear matching during hybridization on a genome-wide scale and the ability to detect these interactions with commonly used approaches that partition nuclear genes into classes based on their role in mitochondrial function. We categorized nuclear genes into three groups based on whether the encoded protein (1) interacts with mitogenomic gene products (interacting n-mt), (2) functions in mitochondria but does not physically interact with mitogenomic gene products (non-interacting n-mt), or (3) does not function in mitochondria (non-n-mt). This *a priori* partitioning allows for an unbiased test of whether selection for matched mitonuclear genotypes leaves genome-wide signatures. Overall, we found inconsistent statistical evidence for differences among these categories based on analyses of mitonuclear interactions in *Xiphophorus* hybrids. However, we found that interacting n-mt genes with larger amounts of non-synonymous substitutions may have more severe consequences for mitonuclear incompatibilities. Therefore, the role of n-mt genes may not be generally detectable on a genome-wide scale, and the sensitivity of these tests should be taken into consideration when interpreting results.

## METHODS

### Genomic data from *Xiphophorus* hybrid populations

Three datasets resulting from previous local ancestry inference in hybrid *Xiphophorus* fishes were acquired as described previously [37,40,43]. Each dataset includes samples from a population of hybrid *Xiphophorus* individuals inhabiting sections of rivers in the states of Hidalgo or San Luis Potosí, Mexico. The first (Calnali Low, or CALL) is on the Calnali River and consists of *X. birchmanni* and *X. malinche* hybrids. The second is at Chahuaco Falls (CHAF) on the Calnali River. It is separated from the CALL population by a waterfall (∼50m tall) [43] and consists of hybrids of the same pair of species as CALL. The third represents two populations (Huextetitla and Santa Cruz, or HUEX-STAC) downstream of the same fork on the Santa Cruz River and derived from a shared hybridization event between *X. birchmanni* and *X. cortezi* followed by independent evolution [37,38].

These datasets were originally analyzed using the *ancestryinfer* pipeline [45] and resulted in posterior probabilities of species ancestry at between 628,881 and 689,982 ancestry informative markers (AIMs) across the nuclear and mitochondrial genomes (Table 1), as described previously [36,39,45]. Informative sites were defined as those with at least a 98% frequency difference between the parental species, as previously described [33,44]. We used a published annotation of the *X. birchmanni* reference genome [46] to identify genes containing each AIM. We then transformed AIM probabilities into categorical genotype calls based on a posterior probability threshold. If a probability for a parent species was greater than or equal to 0.9 and the other parent species’ probability of ancestry was less than or equal to 0.1, ancestry was assigned to the major probability. If instead the sum of both parents’ ancestries was less than or equal to 0.2, the AIM was assigned as heterozygous. AIMs that did not meet these criteria were filtered out of the dataset. Allele counts were then averaged across all AIMs that passed the prior filtering to calculate an average major parent ancestry proportion per individual. These values were then averaged for nuclear and mitochondrial AIMs to obtain population-level averages (Table 1).

**Table 1.** Summary of Xiphophorus hybrid populations used in this study.

| Site Abbreviation | CALL (Calnali Low) | CHAF (Chahuaco Falls) | HUEX-STAC (Huextetitla and Santa Cruz) |
| --- | --- | --- | --- |
| Population Site | Calnali River | Calnali River | Santa Cruz River |
| Major Parent Species | *X. malinche* | *X. malinche* | *X. cortezi* |
| Minor Parent Species | *X. birchmanni* | *X. birchmanni* | *X. birchmanni* |
| Number of Individuals | 281 | 250 | 255 |
| Mitochondrial Haplotype Frequency<sup>a</sup> | 63.7% *X. birchmanni* | 93.5% *X. malinche* | 97.8% *X. cortezi* |
| Average Major Parent Nuclear Ancestry | 52.0% | 71.3% | 88.7% |
| Nuclear Ancestry Informative Sites | 629,430 | 628,788 | 689,966 |
| Mitochondrial Ancestry Informative Sites | 93 | 93 | 16 |
| Estimated Generations Since Initial Admixture | 46 ± 1 [40] | 46 ± 1 [40] | 263 [33] |

<sup>a</sup>Haplotype frequencies for each population were calculated following filtering of mitochondrial AIMs with low-confidence genotypes (posterior probability between 0.2 and 0.8). Filtering resulted in sample sizes slightly lower than the Number of Individuals row: CALL n = 278, CHAF n = 249, HUEX-STAC n = 243.

### Identification of *Xiphophorus* n-mt genes

The gffread [47] program was used to obtain predicted protein sequences, using the -V option to remove any coding sequence with an in-frame stop codon. To annotate n-mt proteins, we compared the *Xiphophorus* translated proteome to UniProt human (UP000005640, GCA_000001405.27) and mouse (UP000000589, GCA_000001635.8) protein sequences. N-mt proteins have been empirically identified in humans and mice as part of the MitoCarta 3.0 project [4]. Because the UniProt human and mouse datasets differed in nomenclature from the MitoCarta dataset, we assigned protein names by searching for MitoCarta sequences in the UniProt human sequences with BLASTP v2.15.0+. A custom Perl script was then used to parse the BLASTP output, and identified proteins were manually confirmed to be the correct protein if percent coverage and sequence identity were not 100%. OrthoFinder v2.5.4 [48] was then run with the primary translated transcript for each *Xiphophorus* sequence and the full sets of human and mouse protein sequences.

MitoCarta protein classifications were used to differentiate non-n-mt from n-mt orthogroups. Interacting and non-interacting orthogroups were distinguished based on a previously published [49] list, updated here, of interacting n-mt proteins that includes OXPHOS subunits, ribosomal subunits, aminoacyl-tRNA synthetases, and DNA/RNA polymerases (Table S2).

### Linkage disequilibrium calculation

We used mitonuclear linkage disequilibrium (LD) as a metric for the association between mitochondrial and nuclear alleles in the CALL and CHAF hybrid populations (see below for analysis methods for the HUEX-STAC population, which differed because their mitogenome is fixed for the haplotype from the *X. cortezi* parental species [38]). Calculations were performed using the *D*′ and *r*^2^ statistics [50]. In the equations that follow, *A* and *B* represent nuclear and mitochondrial loci, respectively, and *pA* and *pB* represent the corresponding allele frequencies. *pAB* is the frequency of the haplotype *AB* across the two sites. The *A* and *B* alleles were assigned to be from the same parental population so that positive and negative values of LD indicate a higher frequency of parental or recombinant allele combinations (i.e., matched and mismatched), respectively.

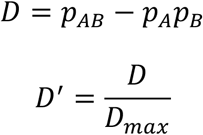

where

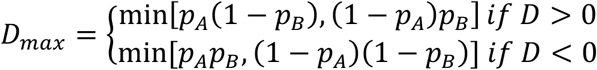

and

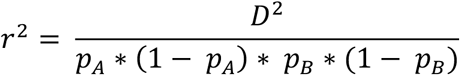

Because all mitochondrial genes are linked on the same non-recombining mitogenome, a single AIM was used (CALL and CHAF’s AIM was in *COX2* and HUEX-STAC’s AIM was in *ATP6*, positions 45 and 846 of GenBank accession CM071424.1, respectively) to identify the mitochondrial haplotype for each fish and to represent the mitogenome in all LD analyses. This approach was appropriate for our data, as all mtDNA AIMs within an individual had approximately the same posterior probability for ancestry assignment. Posterior probabilities were summed across each nuclear AIM as a proxy for frequency of the two alleles. The two LD statistics were calculated for each gene class separately in a custom R script. LD values were visualized using ggplot2 v3.4.0 and ggridges v0.5.4 in RStudio v2023.03.1+446.

LD values were then averaged across AIMs within each nuclear gene to limit pseudo-replication. A one-way ANOVA from the R stats v4.2.2 package (commands *lm* and *anova*) was used to determine whether gene class had a statistically significant impact on LD values. Summary statistics were collected using commands *n*, *mean*, and *sd* from the dplyr v1.1.0, base v4.2.2, and stats v4.2.2 packages, respectively.

In the case of selection on a small number of n-mt genes, shifts in genome-wide averages may not be detectable but could still result in individual loci with the strongest association with the mitogenome being enriched for interacting n-mt genes. We therefore also identified the AIMs and genes with the top 1% of LD values with a custom R script.

### Partial correlation calculation

High LD values can result from population structure (i.e., proportion of parent species ancestry in an individual genome), and the populations used here vary in the proportion of background ancestry (Table 1). We therefore performed a partial correlation (pcor) analysis, which calculated the correlation between mitochondrial and nuclear ancestry while accounting for the correlation expected given the variation in genome-wide ancestry among individuals. AIMs were filtered as described above, and individual ancestry genotype calls for mitochondrial and nuclear AIMs were used to estimate a correlation coefficient with the *pcor.test* function from the R ppcor v1.1 package. This estimate was then averaged across all AIMs in a gene, and a one-way ANOVA was run to determine the effect of gene class on the correlation estimate.

### HUEX-STAC allele frequency calculation

The HUEX-STAC populations are both fixed for the *X. cortezi* mitochondrial haplotype, so we expected *X. cortezi* ancestry to be disproportionately represented at n-mt loci. However, due to the lack of mtDNA variation, the average allele frequency for each nuclear gene was calculated and visualized (with a custom R script) instead of mitonuclear LD to infer how mtDNA background affects selection on n-mt genes in this system. A one-way ANOVA and summary statistics were computed as described above.

A permutation test was then performed to determine whether significant results were robust to violations of normality assumptions in ANOVA. Allele frequency values were permutated by randomly reassigning them across genes 9999 times using base R v4.2.2 functions in a custom R script. Permuted datasets were then used to calculate null expectations for differences in allele frequencies between interacting n-mt and non-n-mt gene classes. The observed difference was compared to the permutations to determine the probability of the pattern arising by chance.

### Non-synonymous substitution analysis

We further filtered the three datasets to include only genes that had substitutions in the coding sequence. We quantified the number of non-synonymous substitutions normalized to the length of the coding sequence. This filtered dataset was analyzed using a linear model with mitonuclear LD as the response variable and an interaction between gene class and our normalized count of non-synonymous substitutions as the explanatory variables. A type III ANOVA was then run (via *Anova* from the R car v3.1-1 package) on the model to determine statistical significance of our predictor variables followed by use of the *summary* function (from base R) with non-n-mts as the reference group. An influential point analysis was performed on the interacting n-mt gene class using DFFIT with the plotting command *ols_plot_dffits* from the R olsrr v0.6.1 package. Identified points were then individually removed from the model with all gene classes to determine their effects on the relationship between mitonuclear LD and non-synonymous substitutions.

### Data and code availability

All code has been deposited in GitHub at https://github.com/sakuster/sensitivity-of-genome-wide-mtnuc. Data can be accessed on figshare (https://doi.org/10.6084/m9.figshare.c.7902011).

## RESULTS

### Identification of *Xiphophorus* n-mt sequences

Of the 24,269 *Xiphophorus* annotated genes, 86.5% (20,988) were assigned to an orthogroup by OrthoFinder. We found that 75.5% (13,264) of the 17,566 identified orthogroups contained at least one *Xiphophorus* gene, and 69.4% (12,198) included at least one gene from all three species (*X. birchmanni*, mouse, and human). Gene and/or genome duplications during the evolutionary history of these three species resulted in some orthogroups with multiple *X. birchmanni* genes (24.1% of orthogroups, or 4,234, contained 2 or more).

These orthogroups were used to classify *Xiphophorus* genes into our three functional categories for each population (Table S3). MitoCarta 3.0 consists of 1,136 human and 1,140 mouse n-mt genes. All but three of the mouse n-mt genes (*Gm4984*, *Htd2*, and *Nsun4*) were identified, and only one human n-mt gene (*NDUFB1*) was not identified in *X. birchmanni*. Overall, 159 orthogroups were found to contain interacting n-mts, 822 to contain non-interacting n-mts, and 15,545 to contain non-n-mts. These orthogroups had 184, 1,042, and 23,043 *X. birchmanni* genes, respectively.

### Mitonuclear linkage disequilibrium does not differ among gene classes in hybrid populations with segregating mitochondrial haplotypes

We sought to determine whether n-mt genes maintained higher mitonuclear LD in *Xiphophorus* hybrids due to selection against mitonuclear incompatibilities. On average, mitonuclear LD values were high across the entire nuclear genome in the CALL and CHAF hybrid populations (Figure 1, Supplemental Figure 1), meaning nuclear ancestry matched the mitochondrial haplotype (positive *D*′ values). This result is also illustrated by differences in average nuclear ancestry between individuals with mitochondrial haplotypes from the two possible parent species (Supplemental Figure 2). These strong associations are likely driven by population structure in these two populations [33,42]. Accordingly, a partial correlation (pcor) test that controls for population structure resulted in lower mitonuclear association values (Supplemental Figure 1).

**Figure 1.**
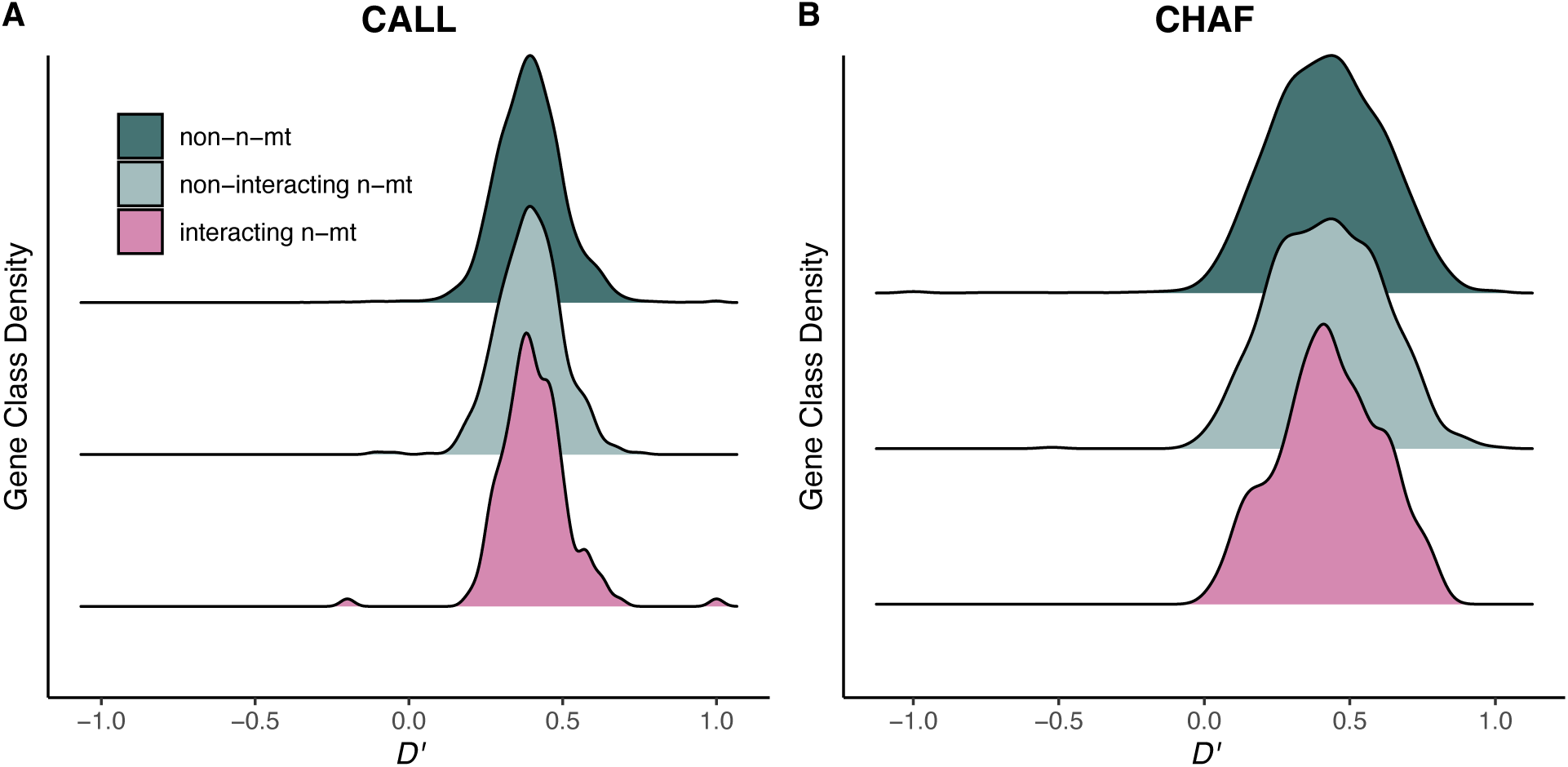
Distribution of mitonuclear D′ values for each gene class in the CALL (A) and CHAF (B) populations. One-way ANOVA testing if D′ value differs among gene classes: CALL p = 0.44 (A); CHAF p = 0.86 (B). Summary statistics are reported in Table S4, and other mitonuclear LD metrics are shown in Figure S1.

Under a model of mitonuclear coevolution, we predicted that the interacting n-mt gene class would have higher mitonuclear LD values compared to the other gene classes. However, we did not find a consistent significant difference between the gene classes for CALL (*D′*: *p* = 0.44; *r*^2^: *p* = 0.44, pcor: *p* = 0.04) or CHAF (*D′*: *p* = 0.86; *r*^2^: *p* = 0.29, pcor: *p* = 0.14; Figure 1, Supplementary Figure 1, summary statistics in Table S4). In fact, the pcor values in CALL instead showed higher mitonuclear LD in the non-n-mt gene class than in either of the two n-mt gene classes. Therefore, it does not appear that mitochondrial haplotypes have preferentially maintained associations with n-mt loci of the same ancestry at a genome-wide scale in these hybridizing populations.

We additionally tested whether interacting n-mt genes were overrepresented in the top 1% of all genes based on our association metrics (*D′*, *r*^2^, pcor). However, again, significant enrichment for interacting n-mt genes was inconsistent (Table S5).

### Known mitonuclear incompatibility genes show positive associations with mitochondrial haplotype

Nine of the 10 previously identified mitonuclear incompatibility genes [44] were included in our CALL analysis, and only two (*NDUFS5* and *SMIM8*) were outliers for mitonuclear LD (with a Z-score of >2 above the mean *D*′ value) (Table S6, Figure S3). Therefore, although the top 1% genome-wide LD values did not show an overrepresentation of n-mt genes, we do find that some genes of known biological importance exhibit signatures of elevated mitonuclear LD. Five other genes (*NDUFA13*, *MTERF4*, *ATP5MG*, *LYRM2* and *RMDN3*) were also above the mean but not outliers by this definition, and the final two genes (*MMUT* and *UQCRC2*) were very close to the mean. Repeating these analyses with *r*^2^ and pcor values instead of *D*′ values resulted in slight changes, but the results were generally similar (Table S6, Figures S4-5).

To see whether these genetic incompatibilities showed similar signatures across populations with varying mtDNA frequencies and different parental source species, we also investigated mitonuclear associations for CHAF and HUEX-STAC (Table S6, Figures S6-9. The CALL and CHAF populations, which are hybrids between the same parental species, exhibit a significant correlation in mitonuclear LD (*D*′) across this set of incompatibility genes (Pearson correlation; *r* = 0.87; *p* = 0.0051). Comparisons with allele frequencies in HUEX-STAC, which originates from a different pair of parental species, also show a positive trend, but the relationships are weaker and/or non-significant with both CALL (*r* = 0.34; *p* = 0.41) and CHAF (*r* = 0.70; *p* = 0.051).

### Interacting n-mt genes show enriched mitochondrial match in an older hybrid population fixed for a mitochondrial haplotype

The HUEX-STAC populations are fixed for the *X. cortezi* mitochondrial haplotype, so we expected to see *X. cortezi* ancestry disproportionately represented at n-mt loci. HUEX-STAC individuals generally have a large percentage of *X. cortezi* ancestry across their entire nuclear genome with a mean allele frequency of 88.7% ± a standard error of 0.011% (Figure 2B). Non-interacting n-mt and non-n-mt genes showed similar mean *X. cortezi* allele frequencies at 82.2 ± 0.02% and 83.1 ± 0.001%, respectively. Interacting n-mt genes had a higher mean allele frequency (88.0 ± 0.097%, Figure 2A, Table S4). Thus, unlike in the CALL and CHAF populations, individuals collected from the HUEX-STAC population did demonstrate a difference in allele frequencies among gene classes (one way ANOVA: *p* = 0.005) and supported the predicted higher degree of matched mitonuclear ancestries for interacting n-mt genes (Figure 2A). This result was also supported by a non-parametric permutation test (Figure S10).

**Figure 2.**
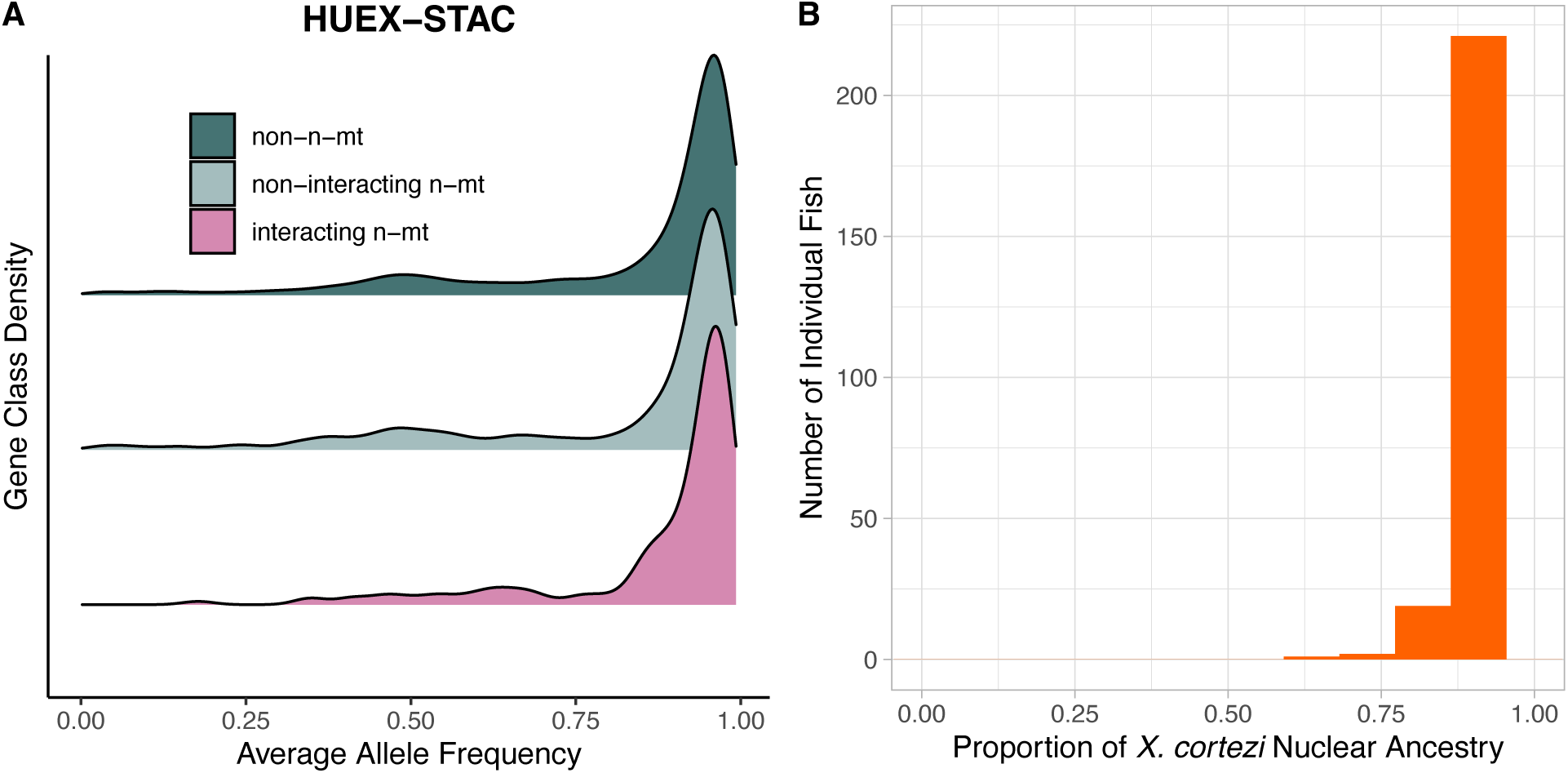
HUEX-STAC analyses for A) allele frequencies for each gene class and B) nuclear genome-wide ancestry for the HUEX-STAC population, a hybrid population fixed for the X. cortezi mitogenome. A one-way ANOVA found gene class to significantly affect allele frequency (p = 0.006); see Table S4 for summary statistics showing differences in means Figure S10 for permutation test demonstrating the robustness of this result. The stacked bar histogram in B was made with 12 bins.

### Interacting n-mt genes with more non-synonymous substitutions may show higher mitonuclear LD

Because many nuclear-encoded proteins may show few or no fixed differences in amino acid sequence between these closely related species, we asked whether the presence of non-synonymous substitutions between species affected how gene class related to mitonuclear LD. There were inconsistent effects of gene class, non-synonymous substitutions, and the interaction between these two across metrics and populations (Tables S7-8). There was, however, some evidence that interacting n-mt genes with more non-synonymous substitutions have higher mitonuclear LD compared to non-n-mt genes (Figure 3, Figures S11-12, Table S9). The significant interactions in CALL and CHAF were found to be the result of a few influential genes (Table S10). For example, *NDUFS5*, a gene previously shown to underly a lethal mitonuclear incompatibility, consistently appeared as influential across association metrics and populations where a significant interaction was found. When *NDUFS5* was removed from the dataset, the difference between the interacting n-mt and non-n-mt gene classes was eliminated. Other known incompatibility genes were not consistently influential, although *NDUFA13* and *ATP5MG* were detected in some analyses in CHAF.

**Figure 3.**
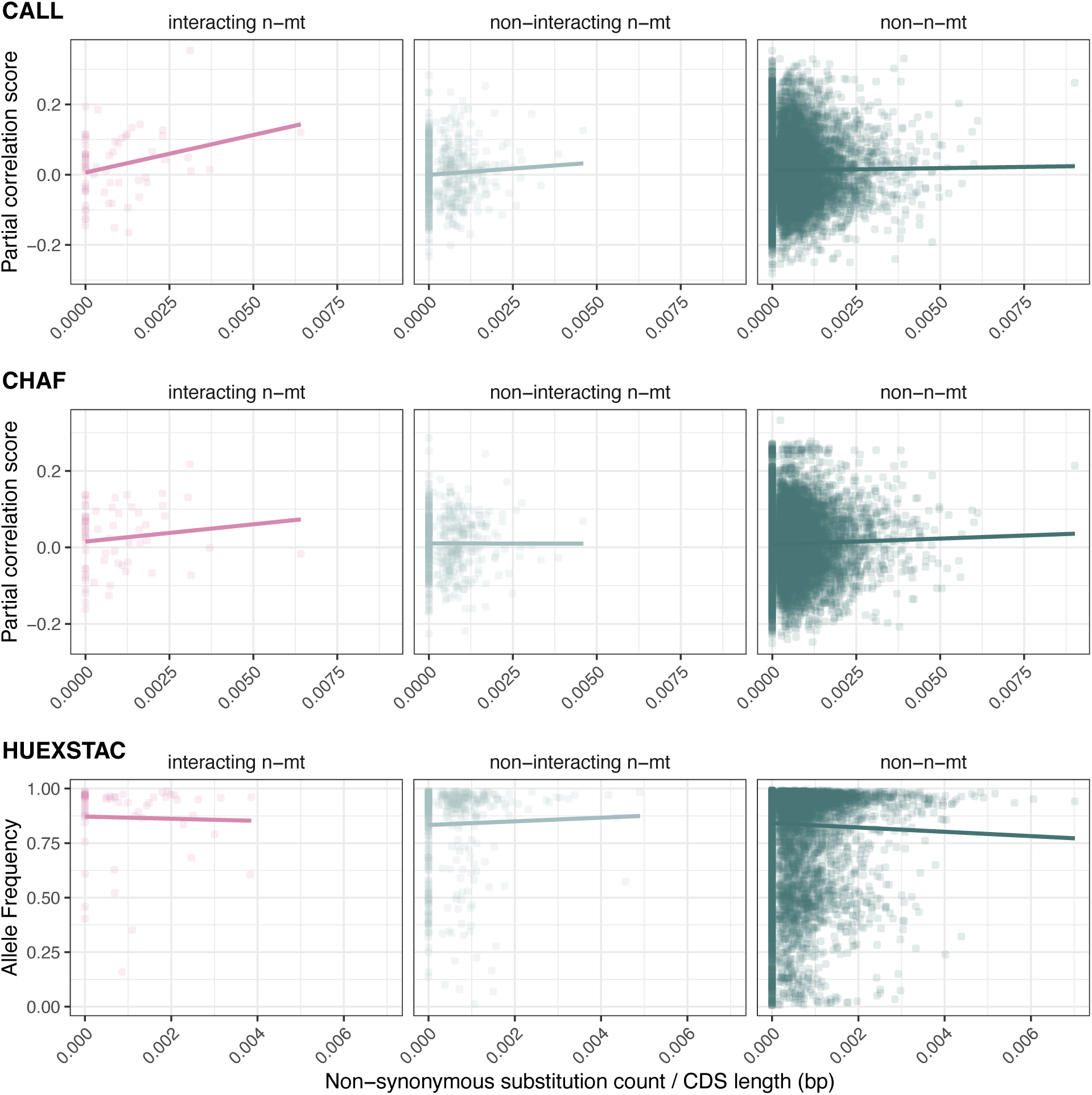
Regression analysis of mitonuclear association metric by non-synonymous substitutions. Partial correlation values for CALL do not show a significant interaction (type III ANOVA p = 0.073) but interacting n-mts have a stronger linear relationship than the non-n-mt relationship (p = 0.034). Partial correlation values for CHAF does not have a significant interaction (p = 0.67) or difference in slopes (p = 0.50). X. cortezi allele frequencies in HUEX-STAC also do not show a significant interaction (p = 0.43) or a difference in slopes (p = 0.85). Other metrics of mitonuclear LD can be found in Figures S10-11. Statistics can be found in Tables S7-10.

## DISCUSSION

We tested the prediction that the interacting n-mt gene class would show the strongest “matched” association with mitochondrial haplotype. Despite the known role of n-mt genes in hybrid incompatibilities between *X. birchmanni* and *X. malinche*, the CALL and CHAF populations did not support this prediction. This negative result likely speaks to the statistical power issues inherent in identifying mitonuclear interactions. The sensitivity of genome-wide tests for signatures of selection acting preferentially on n-mt genes will likely be affected by the structure of the incompatibility and the genomic background of the genes under selection, factors that are also important for detecting loci involved in nuclear-nuclear hybrid incompatibilities [51]. Specifically, the number of genes under selection, where those genes are distributed in the genome, and/or whether only n-mt genes are involved could affect the sensitivity of these tests.

N-mt genes make up a small portion of the genome, and only a subset of those genes may be participating in mitonuclear epistasis. Incompatibilities can potentially involve a few loci with large effect size or many loci of small effect (or some combination of these extremes). The analyses performed by Robles et al. [44] identified 10 n-mt genes among seven genomic regions contributing to mitonuclear incompatibilities in *Xiphophorus* species hybrid populations, one of which was CALL. These genes represent only 6.8% of all interacting n-mt genes and 1.0% of all n-mt genes. Selection on these few genes would have to be exceptionally strong and have had enough time to act to detect a significant shift in the average for the entire interacting n-mt gene class. In contrast, systems such as *Tigriopus* copepods appear to have many genes involved in mitonuclear incompatibilities and have proven to be powerful in advancing the field of mitonuclear genetics and genomics [20,31]. Therefore, even in cases where n-mt genes are contributing to negative mitonuclear epistasis, the number of loci may be too small to detect genome-wide effects with the approach implemented here, which is inherently designed to detect polygenic contributions to incompatibilities.

Elevations in background LD (e.g., due to population structure) and the distribution of n-mt genes across the genome will also affect the ability to detect a difference in mtDNA ancestry matching between n-mt and non-n-mt genes. Any signal of this difference arising from selection on an individual n-mt target could be diluted by hitchhiking at linked non-n-mt loci [52]. This effect would be particularly problematic for populations that are the result of recent admixture events and, therefore, have not had many generations for recombination to break up haplotype blocks in the nuclear genome [41,53], such as in CALL and CHAF (whose initial admixture was estimated to be ∼46 generations ago [35]). In some cases, clustering of n-mt genes in specific regions of the genome could create strong targets of selection that overcome this dilution effect. For example, genomic regions with a high density of n-mt genes have been identified in the long-tailed finch [54] and the eastern yellow robin [55], both of which have found evidence that n-mt genes disproportionately share ancestry with mtDNA.

Additionally, not all mitonuclear incompatibilities are necessarily mediated by n-mt genes. Instead, some incompatibilities may involve nuclear-encoded proteins that are not targeted to mitochondria but nonetheless create genetic interactions with mtDNA through co-functionality in biochemical and signaling pathways that span multiple cellular compartments [56]. For example, our definition of an n-mt gene is a gene whose product is targeted for import into mitochondria. However, mitochondria are a part of a complex cellular network, and any number of cellular processes could affect mitochondrial function. Genes involved in these processes are, by our definition, non-n-mt genes. Therefore, any contributions of non-n-mt genes to mitonuclear incompatibilities would directly undermine search strategies based on partitioning the genome into n-mt and non-n-mt categories.

However, these challenges were somewhat mitigated in our analyses of CALL and CHAF by including the number of non-synonymous substitutions for each gene in the model. By looking only at genes with substitutions between species, we uncovered a pattern in which interacting n-mt genes tended to show a stronger relationship between mitonuclear association and the number of non-synonymous substitutions than non-n-mt genes (Figure 3, Figures S11-12). This stronger, positive relationship could imply that putative functional divergence results in stronger selection to match ancestry to the mitochondrial haplotype if the divergence occurs in an interacting n-mt gene. However, this relationship was found to be disproportionately driven by a few genes, namely *NDUFS5*, a known incompatibility gene [43]. Nevertheless, this result suggests that including protein sequence divergence could increase the power to detect the effects of mitonuclear incompatibilities, even when using genome-wide tests in a system with few genes of large effect. We speculate that these effects will be easiest to detect in interacting n-mt genes but that this result does not rule out the possibility of other genes contributing to mitonuclear incompatibilities.

Despite the challenges outlined above, one of the three hybrid populations we investigated (HUEX-STAC) supported our prediction of enrichment in matched ancestry for n-mt genes. Notably, this hybrid population is older than CALL and CHAF, estimated to have formed ∼250 generations ago [33,35], and it is the only population we investigated that is fixed for a mitochondrial haplotype from just one of the parental species. Its older age suggests there has been more time for incompatible genotype combinations to be purged from the population and for recombination to have broken up haplotype blocks in the nuclear genome, thereby limiting the effects of hitchhiking. We therefore speculate that the age of HUEX-STAC and the fixation of a mitochondrial haplotype may have made selection on n-mt loci stronger and/or more effective. In such populations, half of the possible mitonuclear combinations have been removed, so selection for matched ancestry would consistently favor the same n-mt allele. In contrast, when mitochondrial haplotypes are still segregating, the fitness effects of an n-mt allele may change depending on an individual’s mtDNA. Because mitochondrial and nuclear loci are not physically linked, very strong selection would be needed to preserve mitonuclear LD in segregating populations [57].

Despite the potential importance of the age and fixed mitochondrial haplotype of the HUEX-STAC population, previous studies have found examples of hybrid populations with both fixed and segregating mitochondrial haplotypes that show a disproportionate effect of selection on n-mt genes during hybridization (Table S1). Therefore, expanding the availability of genomic datasets from hybrid systems and performing a formal meta-analysis would be important future directions to assess relevant variables, such as whether mitochondrial haplotypes are segregating in the population, the age of the hybrid population, the number of loci involved and their effect sizes, and the genetic structure of the population. Reanalysis of publicly available genome-level data from hybrid populations may also be a valuable approach to control for method of ancestry inference and metric of association between mtDNA and nuclear loci.

Another variable that differentiates the HUEX-STAC population is the parental makeup of the hybrids. Although geographically distinct, CHAF and CALL share parental species (*X. birchmanni* and *X. malinche*). In contrast, HUEX-STAC was formed by hybridization between *X. birchmanni* and *X. cortezi*. These three species demonstrate mitonuclear discordance [58] and a history of introgression [42], both of which may impact mitonuclear incompatibilities in these hybrids, potentially resulting in stronger selection for matched mitonuclear ancestry. Given these different demographic histories, each of the hybrid populations we analyze here is expected to have numerous unique features, so caution is warranted in attributing any of our results to a single feature (such as the fixed mitochondrial haplotype in HUEX-STAC). Nevertheless, we view the potential role of fixed vs. segregating mitochondrial haplotypes to be an important area of future research given the small number of studies conducted to date.

Overall, our findings suggest that splitting up the genome into n-mt and non-n-mt gene classes may be an underpowered method to detect effects of mitonuclear epistasis in recently admixed populations, even when there are clear biological effects of said epistasis. Although such tests for genome-wide signatures may offer a relatively unbiased approach, they may be susceptible to false negatives. Therefore, important contributions of mitonuclear interactions to speciation may remain unrecognized in many systems.

## ACKNOWLEDGEMENTS

SAK was supported by an NIH T32 fellowship (GM132057) and an NSF GRFP (006784). This work was also supported by NIH grants to DBS (R35GM148134) and JCH (R35GM142836) and an EDGE - NSF grant (IOS-2421661) to JCH and MS. We thank three anonymous reviewers for their insightful comments.

## Supplementary Materials for the manuscript

### SUPPLEMENTAL FIGURES

**Figure S1.**
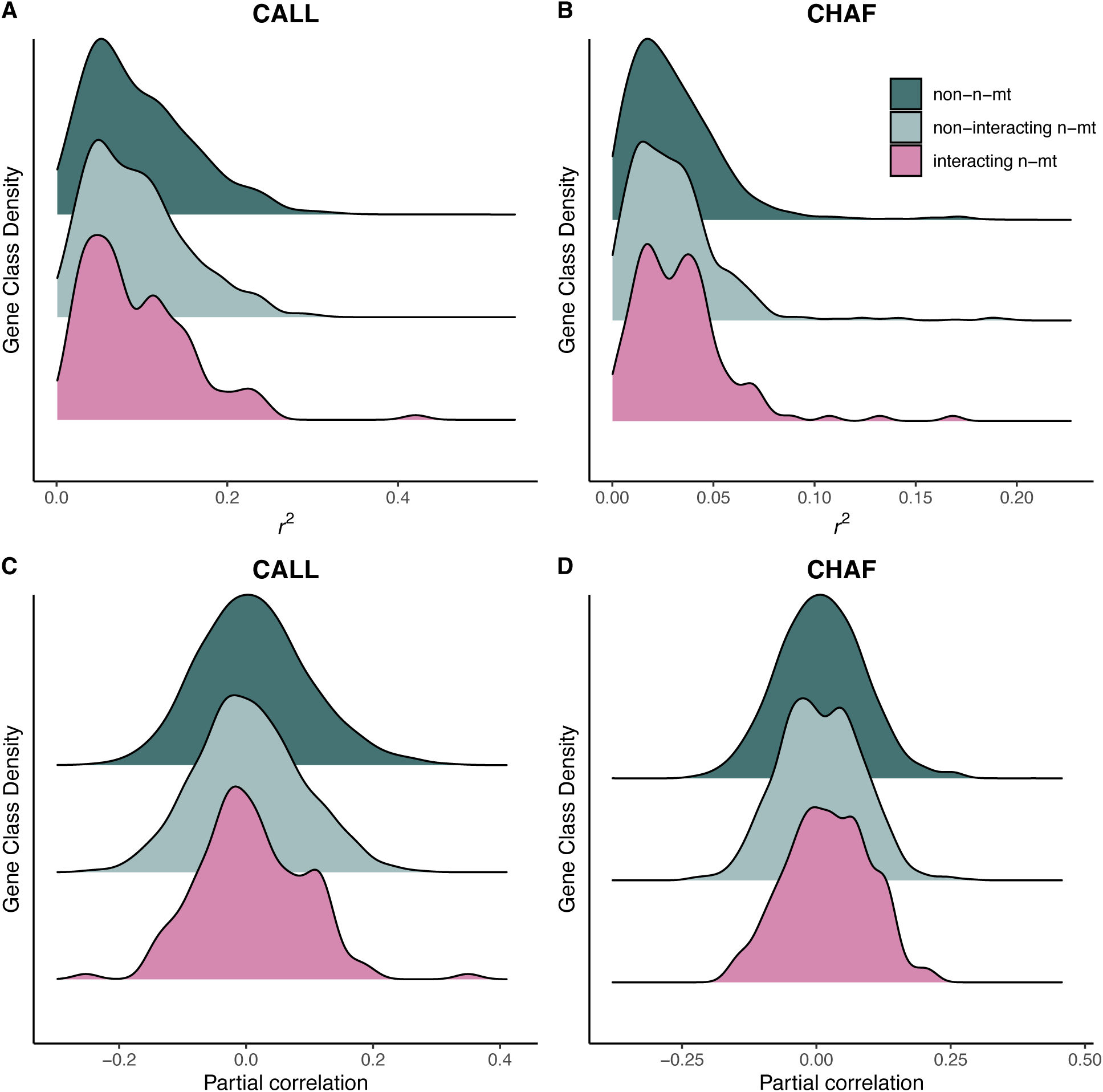
Other metrics of mitonuclear association values for each gene class in the CALL (A and C) and CHAF (B and D) populations. Association is calculated as r^2^ in panels A and B and as a partial correlation that accounts for genome-wide ancestry in panels C and D. Note the change of scale on the x-axis across populations and LD statistics. P-value for a one-way ANOVA testing if gene class affects LD value: CALL r^2^ 0.44 (A), CHAF r^2^ 0.29 (B), CALL partial correlation 0.04 (C), CHAF partial correlation 0.14 (D). Summary statistics are reported in Table S4.

**Figure S2.**
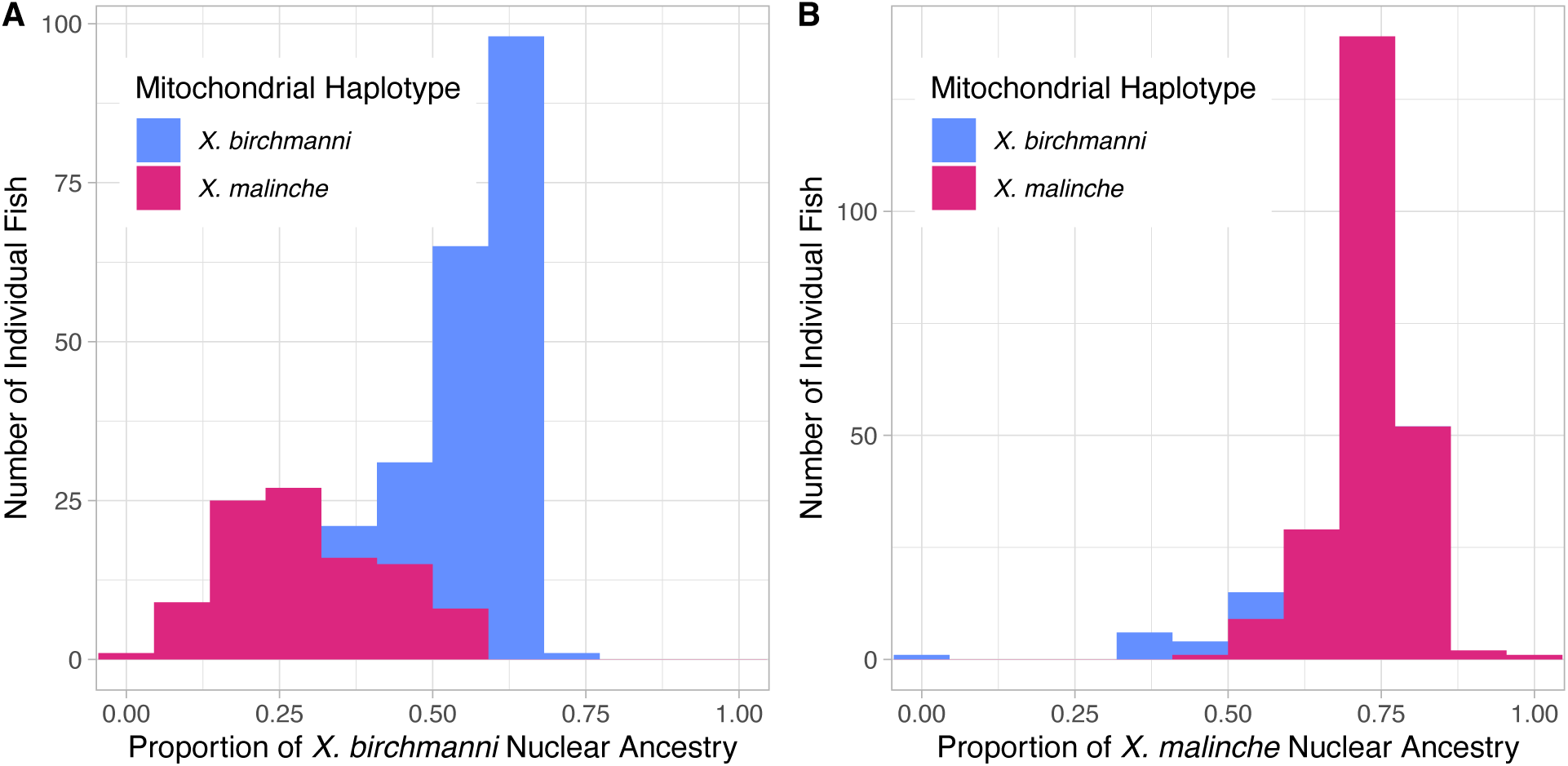
Stacked bar histogram for the proportion of the nuclear genome derived from the major parent for each fish in A) CALL and B) CHAF. Proportion is calculated as the sum of all major parent AIMs divided by number of AIMs. Each fish is colored by which of the two segregating mitochondrial haplotypes it has (blue = X. birchmanni, fuchsia = X. malinche). Both stacked bar histograms were made with 12 bins. Note that y-axis upper limits differ between the panels.

**Figure S3.**
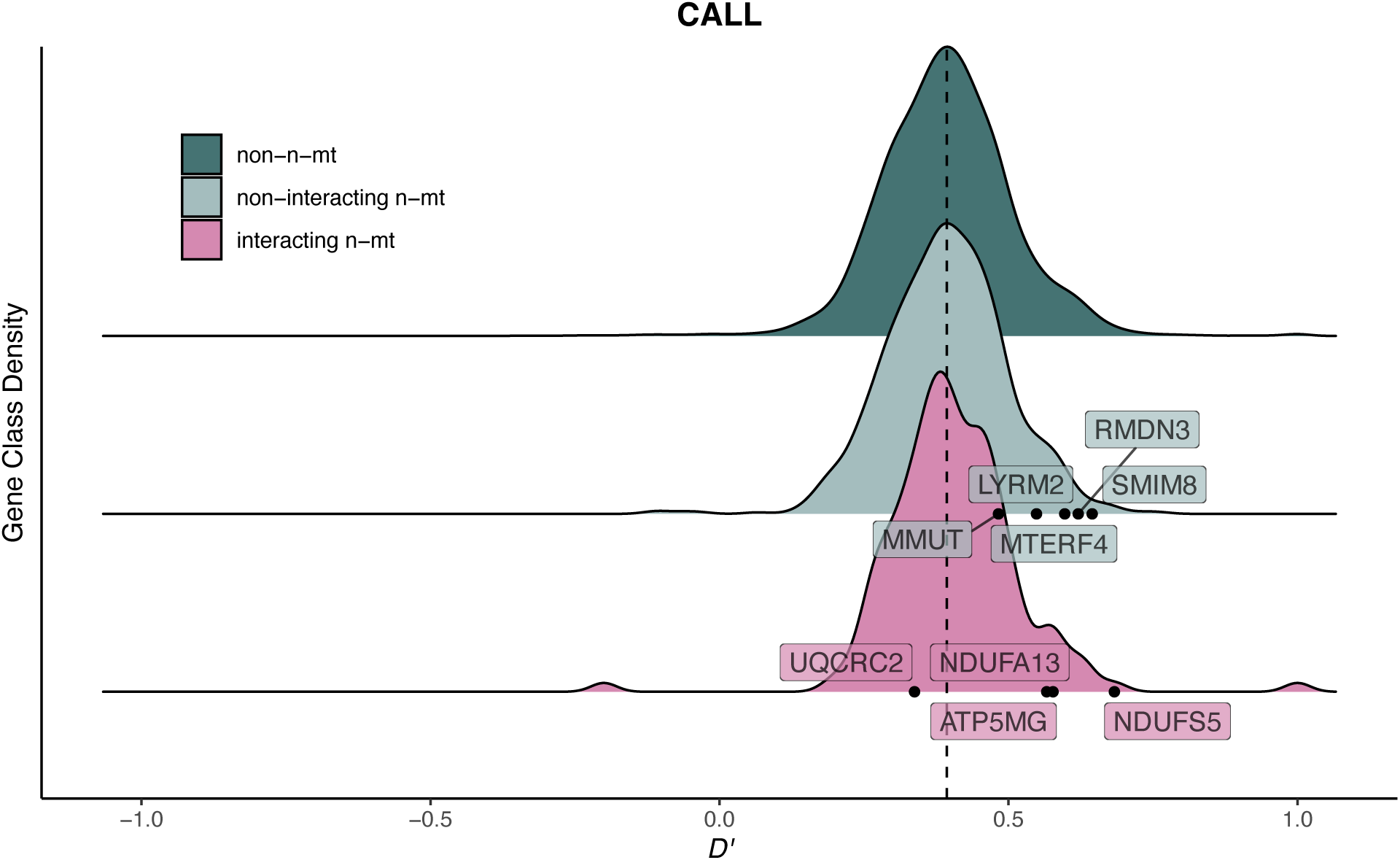
Location of previously identified incompatibility genes [1] in the CALL population D’ values. Dashed line indicates genome-wide mean D’.

**Figure S4.**
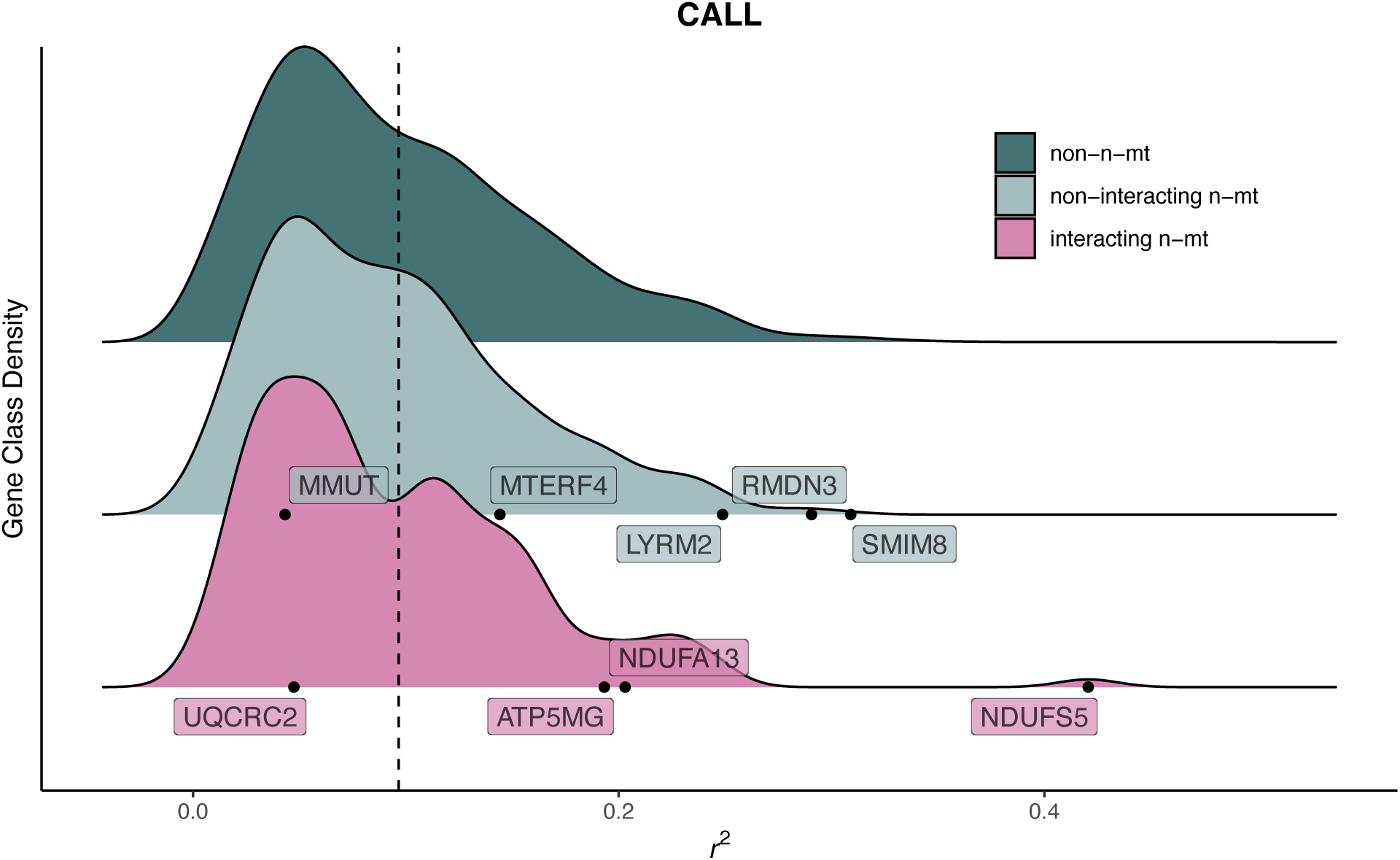
Location of previously identified incompatibility genes [1] in the CALL population r^2^ values. Dashed line indicates genome-wide mean r^2^.

**Figure S5.**
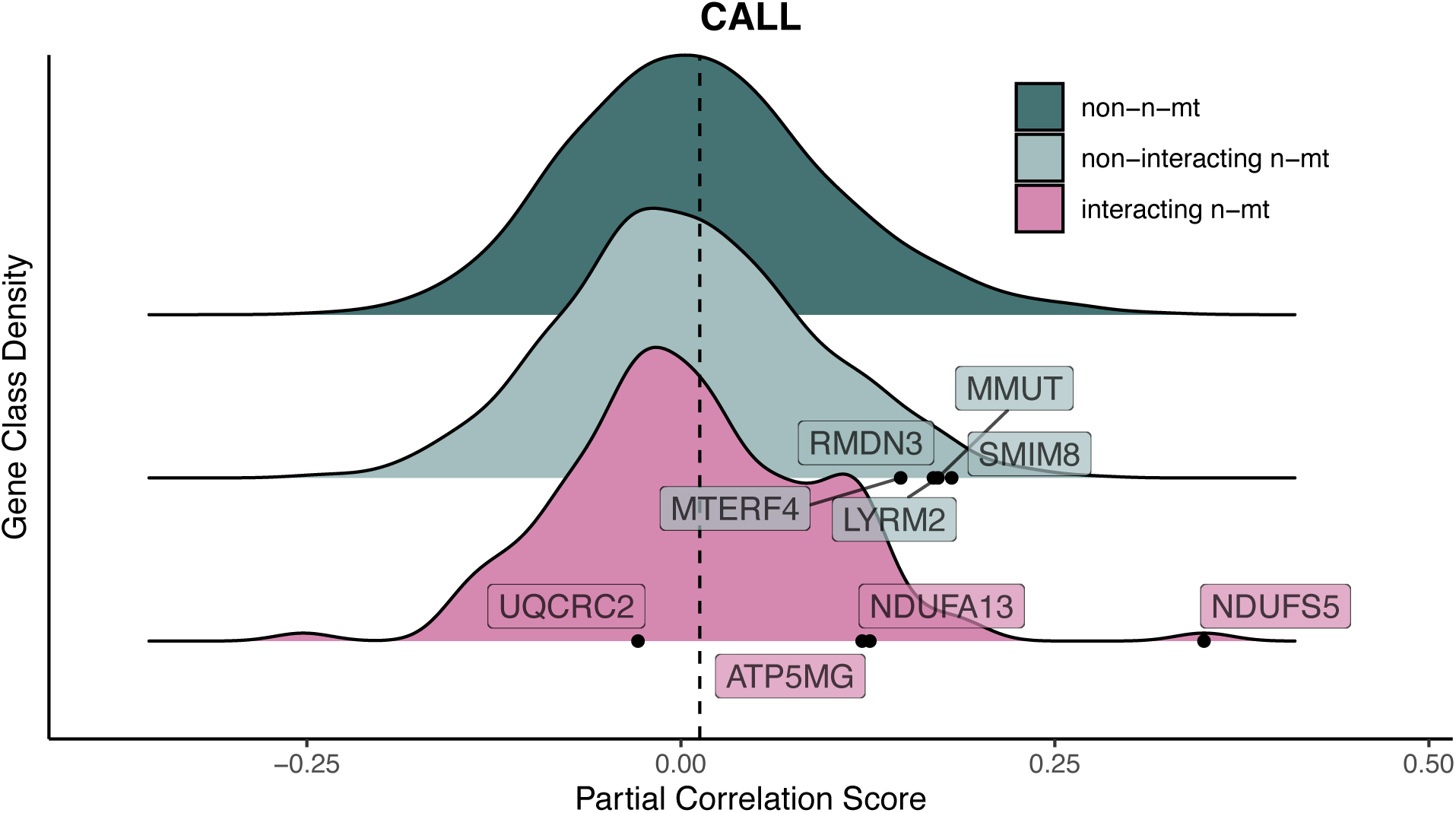
Location of previously identified incompatibility genes [1]in the CALL population partial correlation values. Dashed line indicates genome-wide mean partial correlation score.

**Figure S6.**
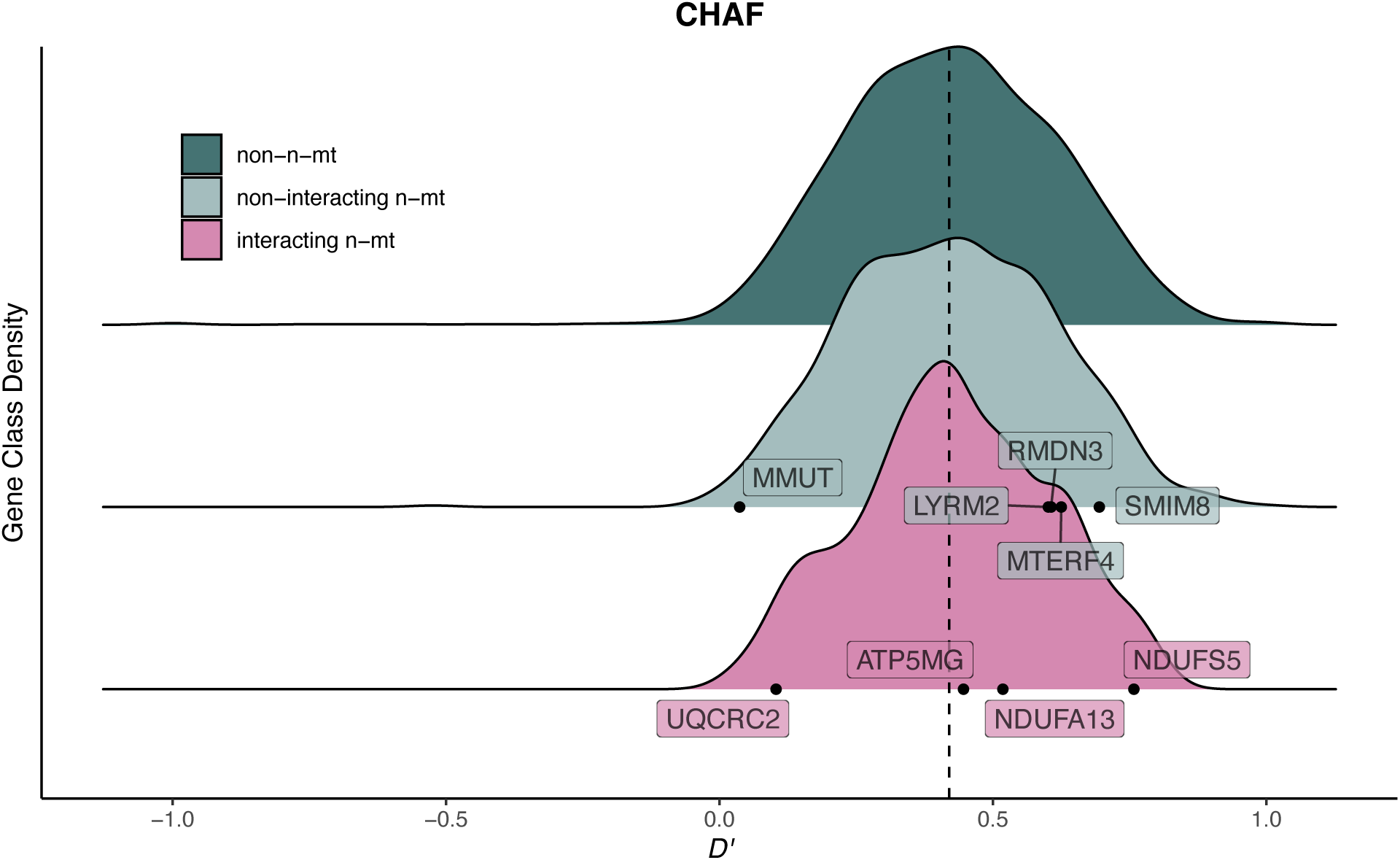
Location of previously identified incompatibility genes [1] in the CHAF population D’ values. Dashed line indicates genome-wide mean D’.

**Figure S7.**
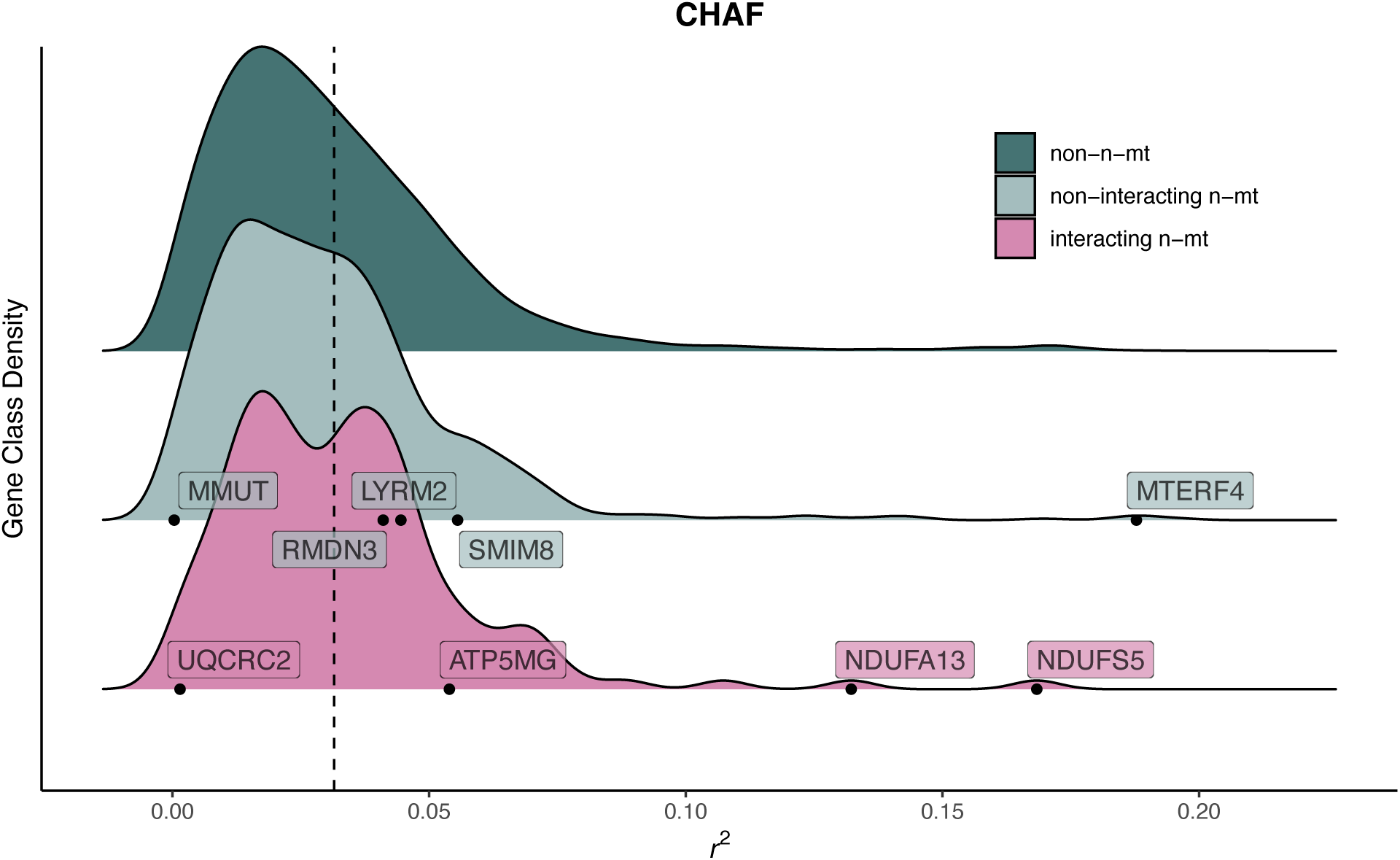
Location of previously identified incompatibility genes [1]in the CHAF population r^2^values. Dashed line indicates genome-wide mean r^2^.

**Figure S8.**
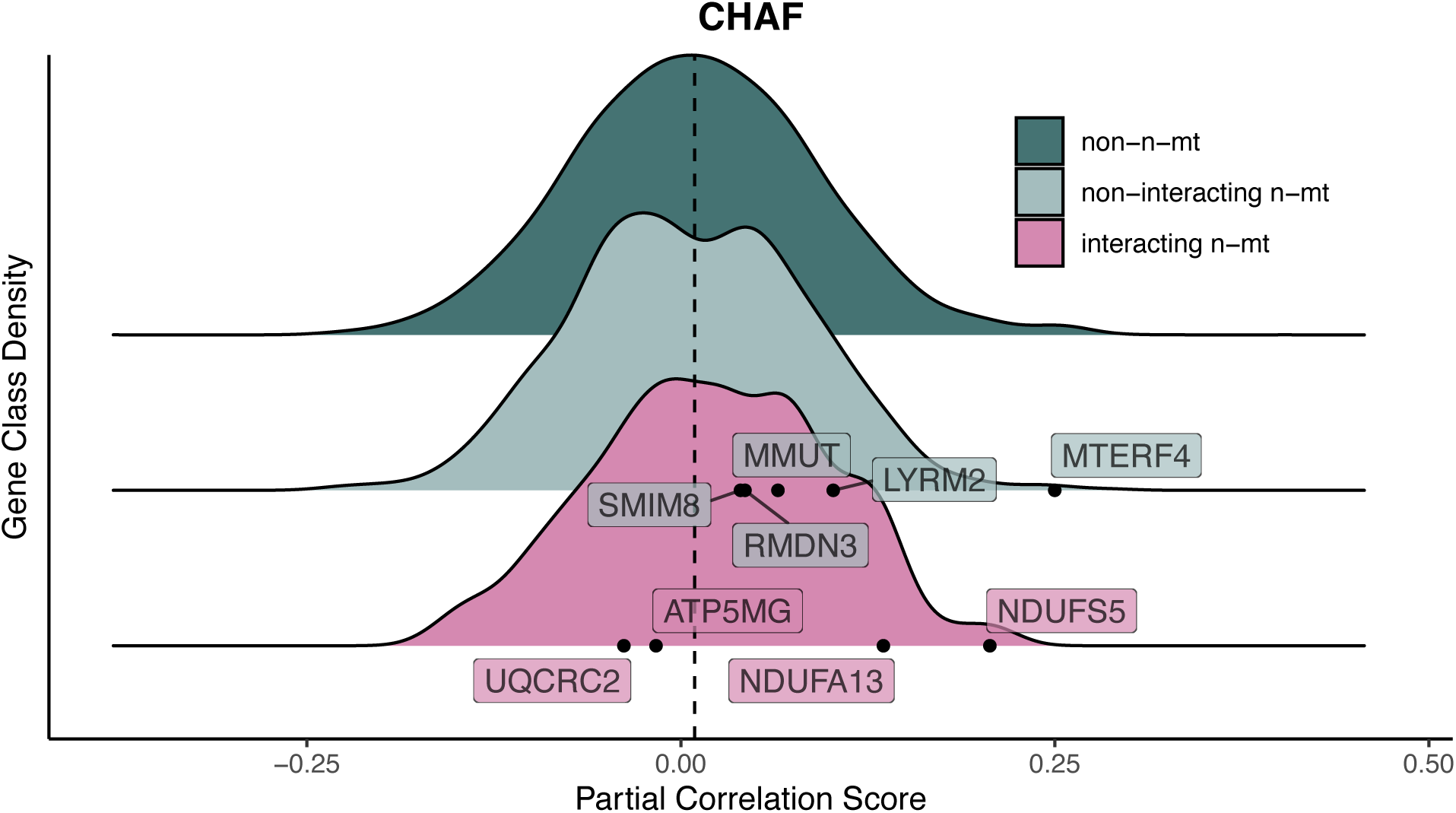
Location of previously identified incompatibility genes [1]in the CHAF population partial correlation values. Dashed line indicates genome-wide mean partial correlation score.

**Figure S9.**
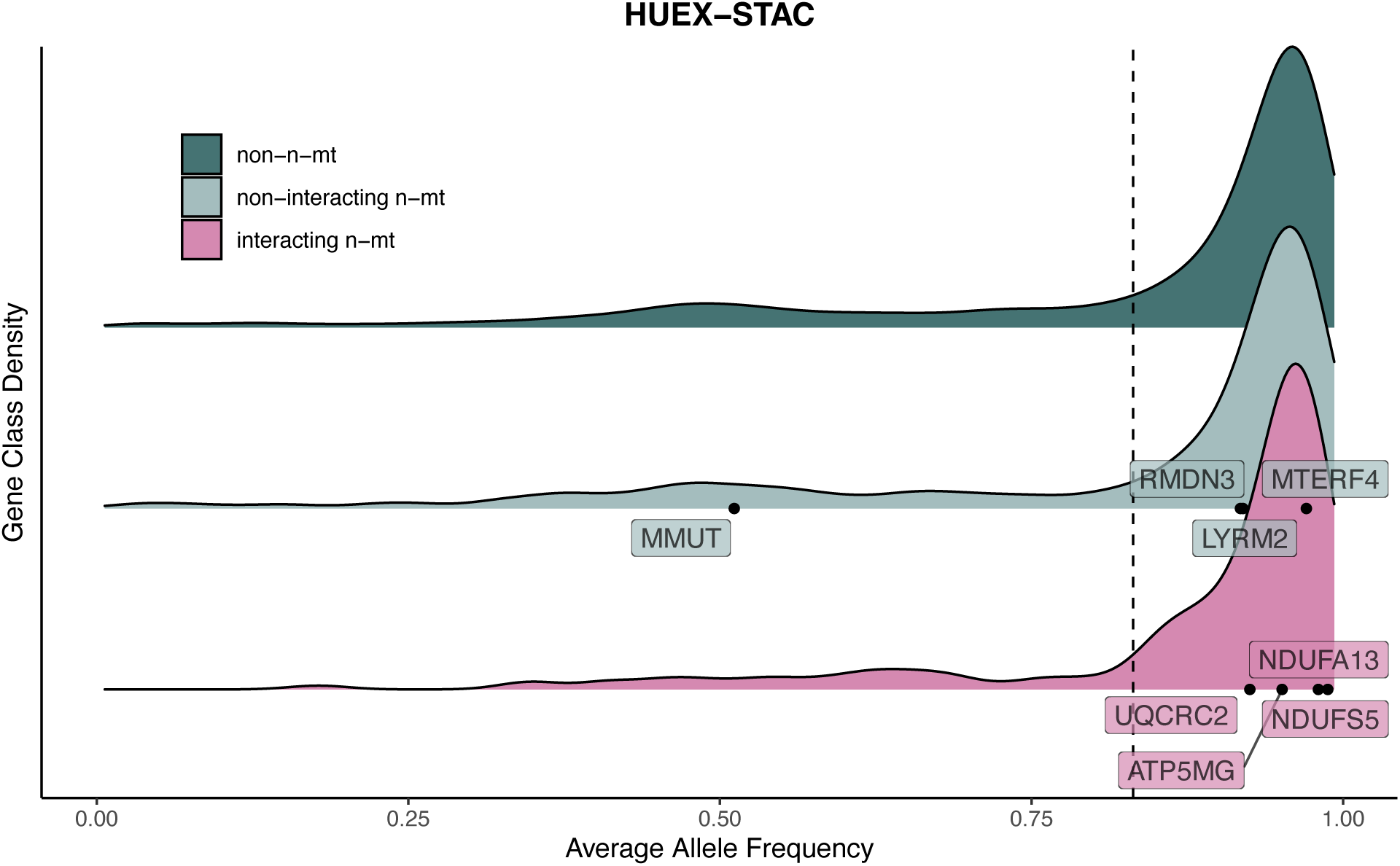
Location of previously identified incompatibility genes [1] in the HUEX-STAC population X. cortezi allele frequency values. Dashed line indicates genome-wide mean allele frequency.

**Figure S10.**
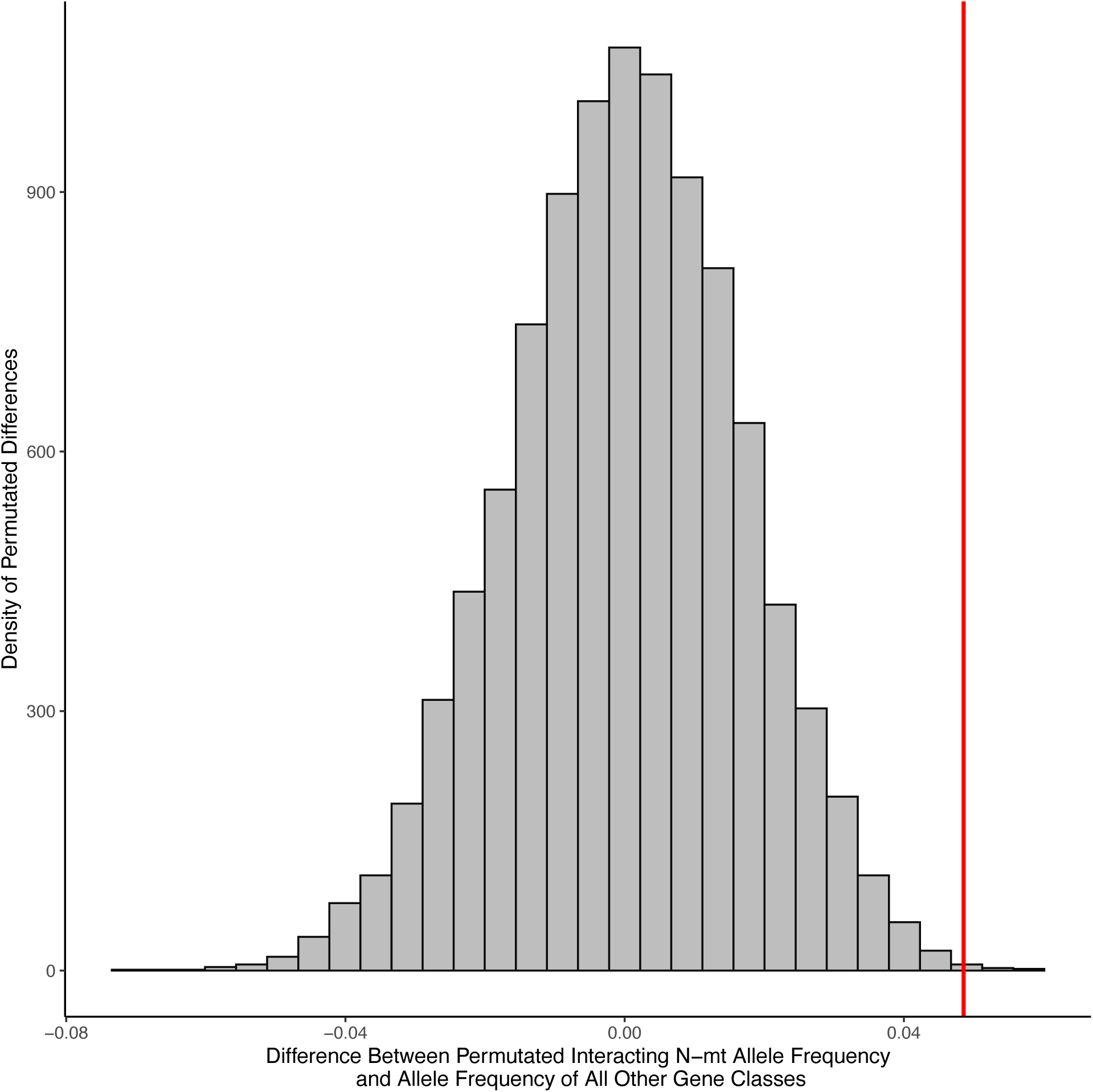
Histogram demonstrating permuted differences in mean allele frequency between all interacting n-mt genes and the mean of all other genes. Red line represents the observed difference in interacting n-mt gene class and all other classes. In this permutation test, we found the interacting n-mt gene class to be different from the other two gene classes (p = 9 × 10^-4^), supporting the parametric analysis (ANOVA) in this population.

**Figure S11.**
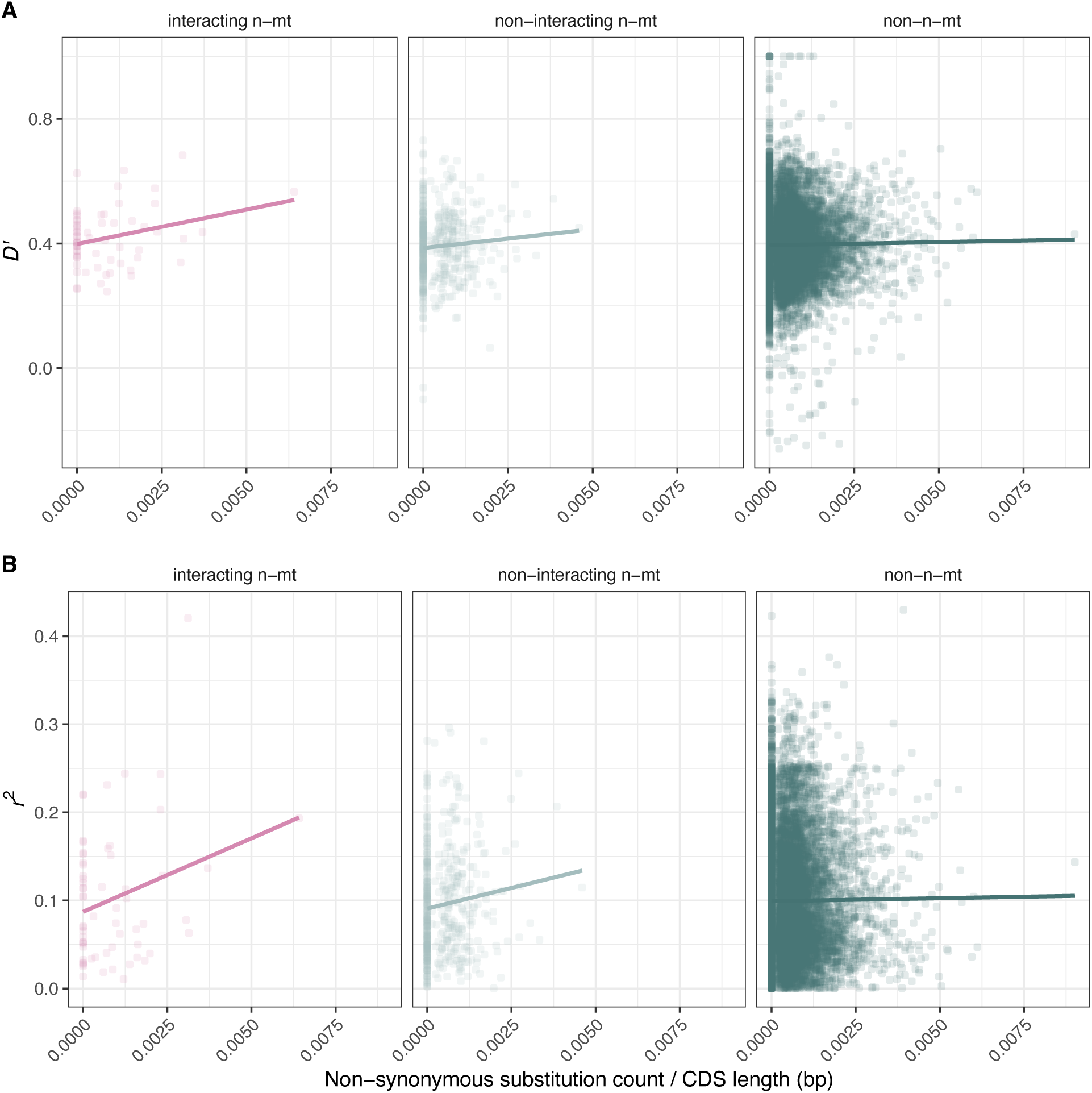
Non-synonymous substitution model for the CALL population for the A) D’ and B) r^2^ metrics. In a model that accounts for an interaction between gene class and non-synonymous substitutions, interacting n-mt genes have a stronger relationship to the r^2^ (p = 0.019) but not the D’ (p = 0.092) metric (Table S9), but see points driving this relationship in Table S10 and summary statistics in Table S7.

**Figure S12.**
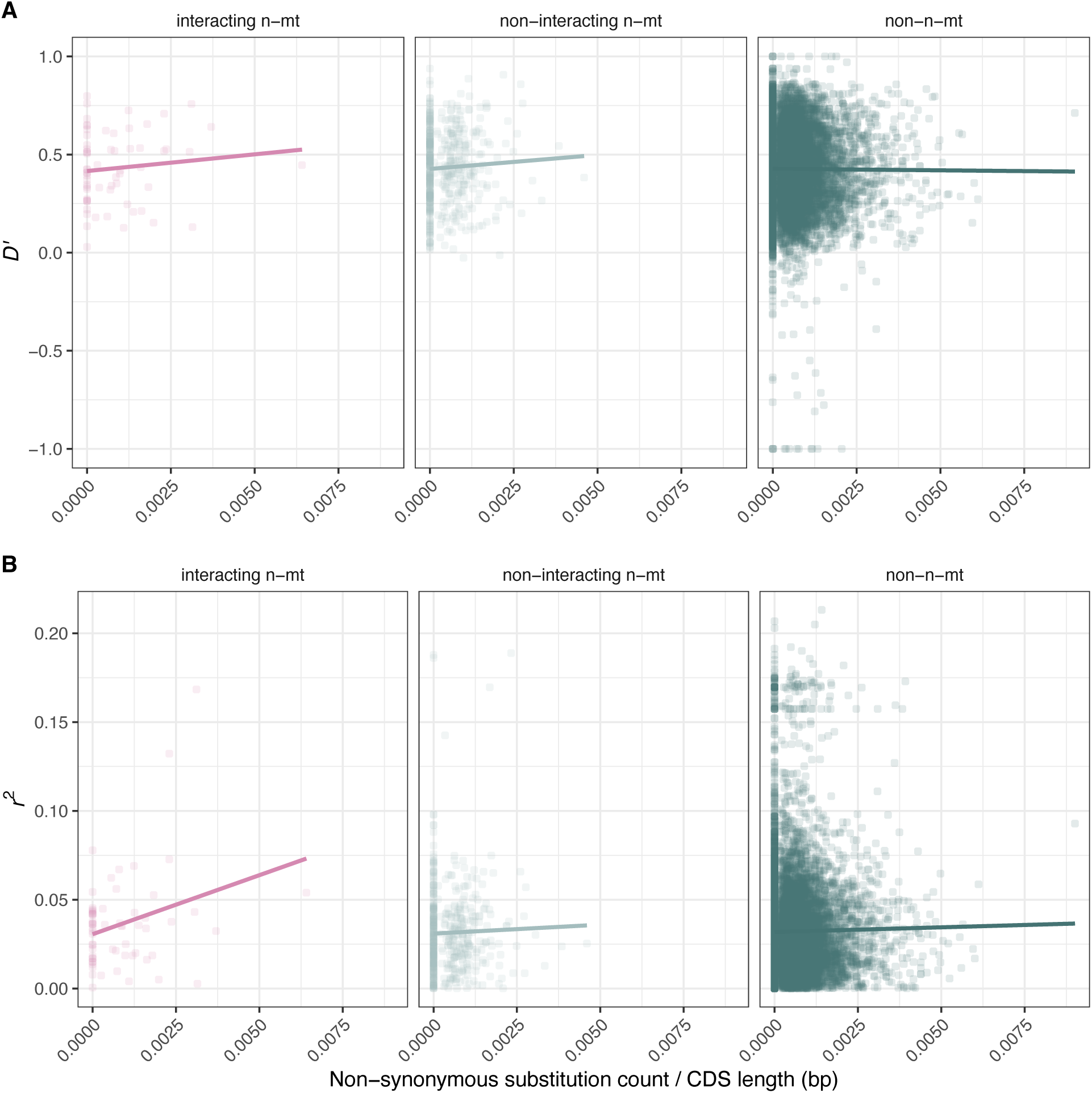
Non-synonymous substitution model for the CHAF. population for the A) D’ and B) r^2^ metrics. In a model that accounts for an interaction between gene class and non-synonymous substitutions, interacting n-mt genes have a stronger relationship to the r^2^ (p = 0.027) but not the D’ (p = 0.37) metric (Table S9), but see points driving this relationship in Table S10 and summary statistics in Table S7.

### SUPPLEMENTAL TABLES

**Table S1.**
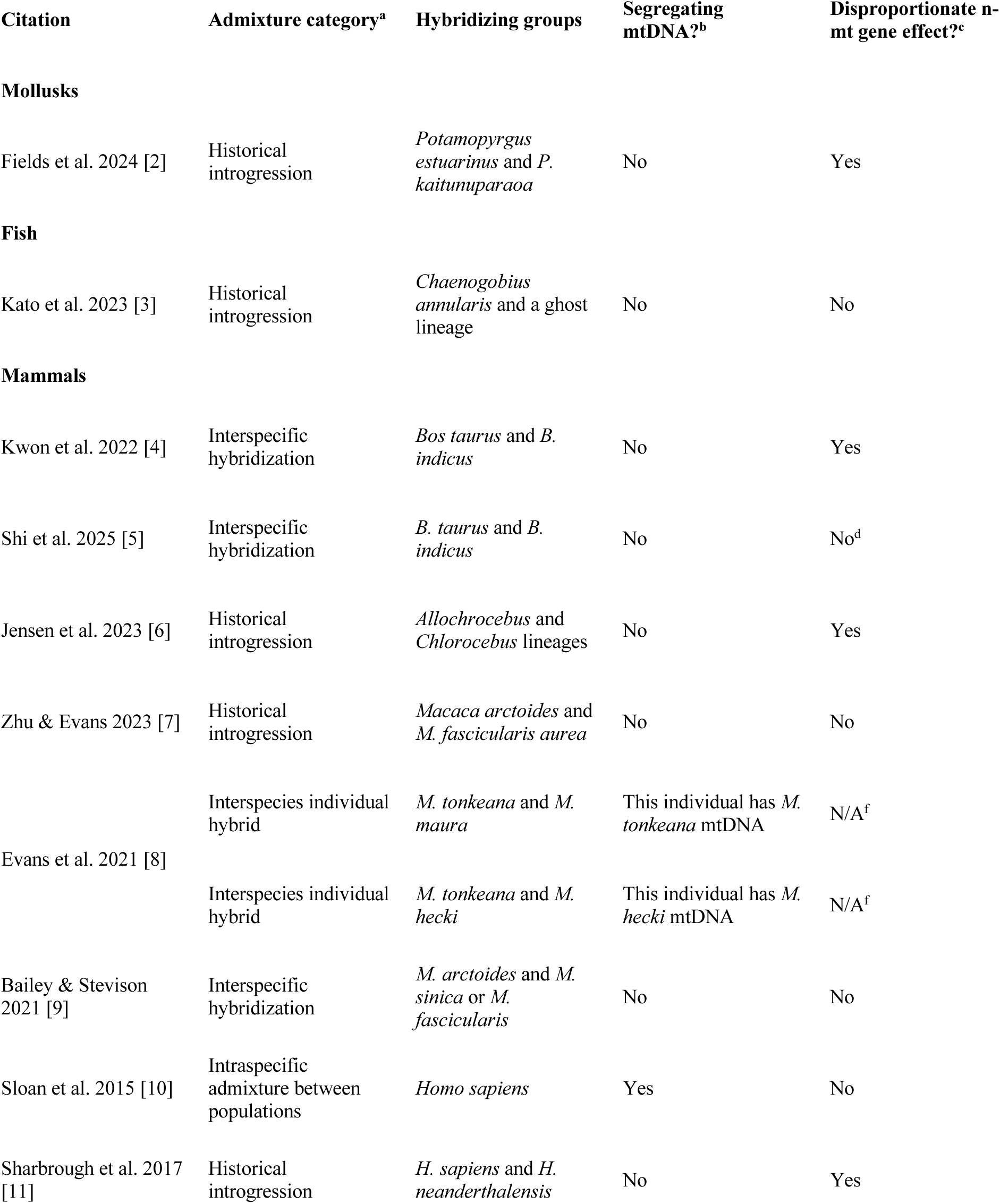
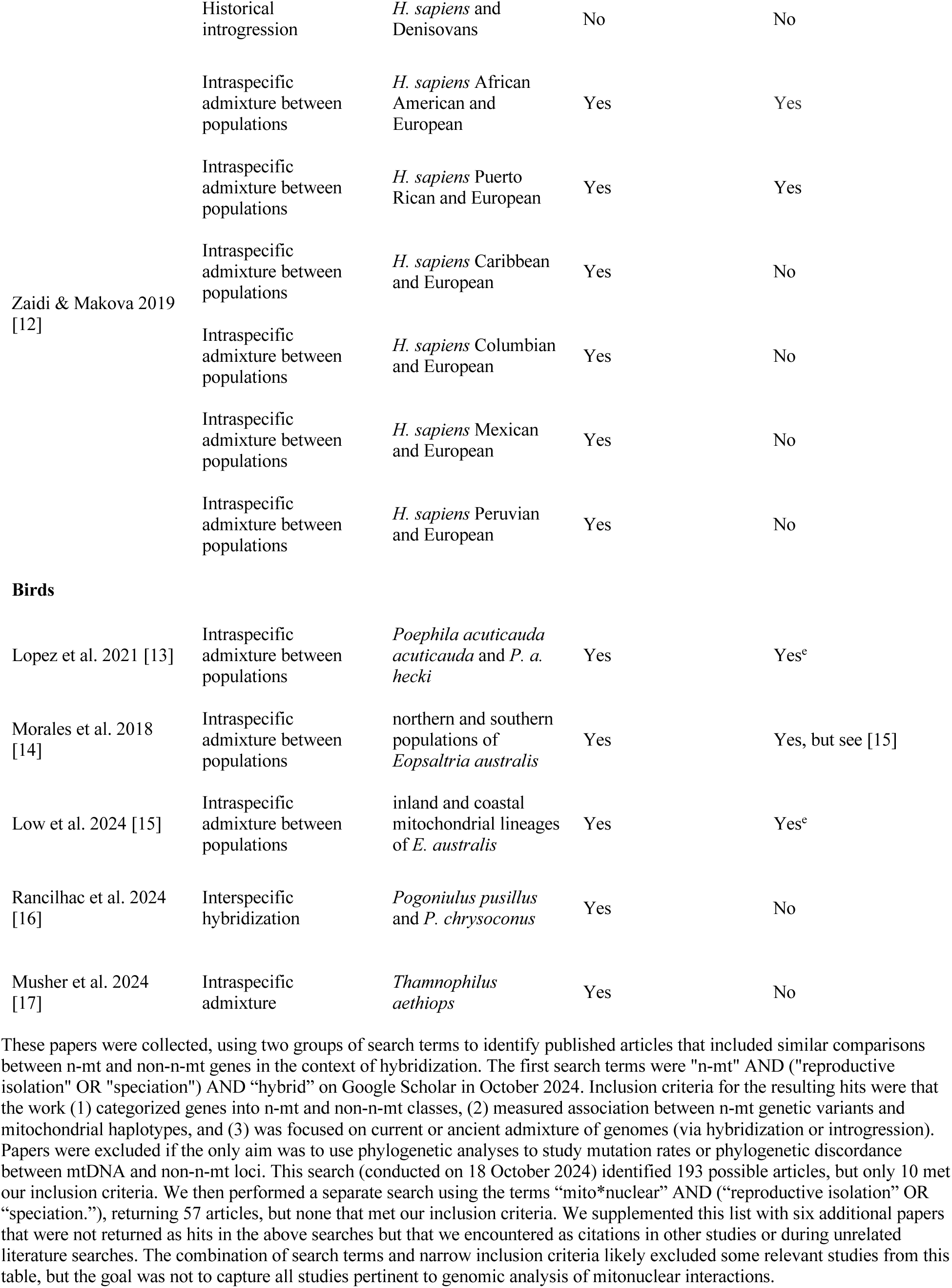
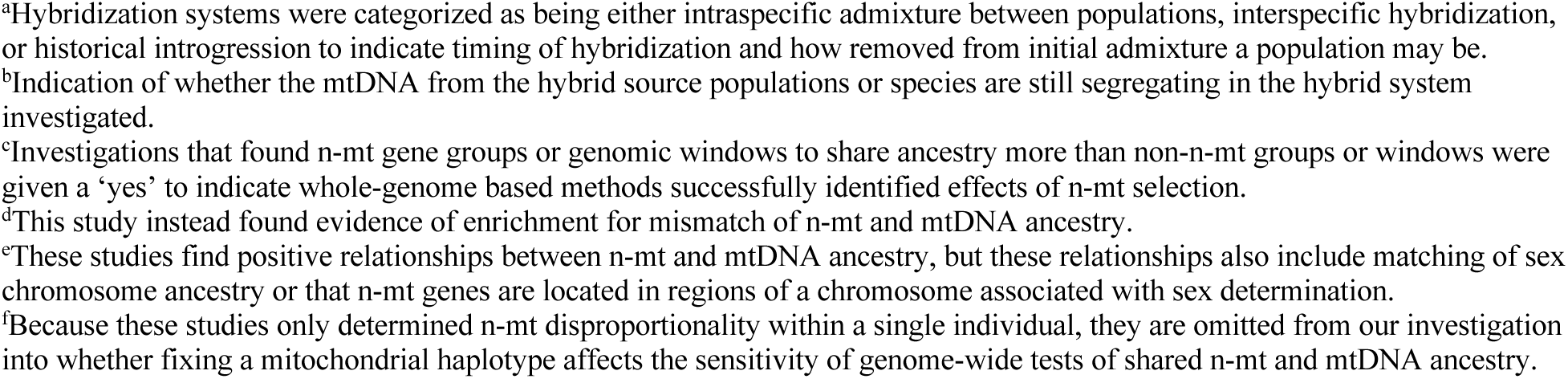
Studies investigating differential effects of n-mt and non-n-mt gene groups or genomic windows.

**Table S2.**
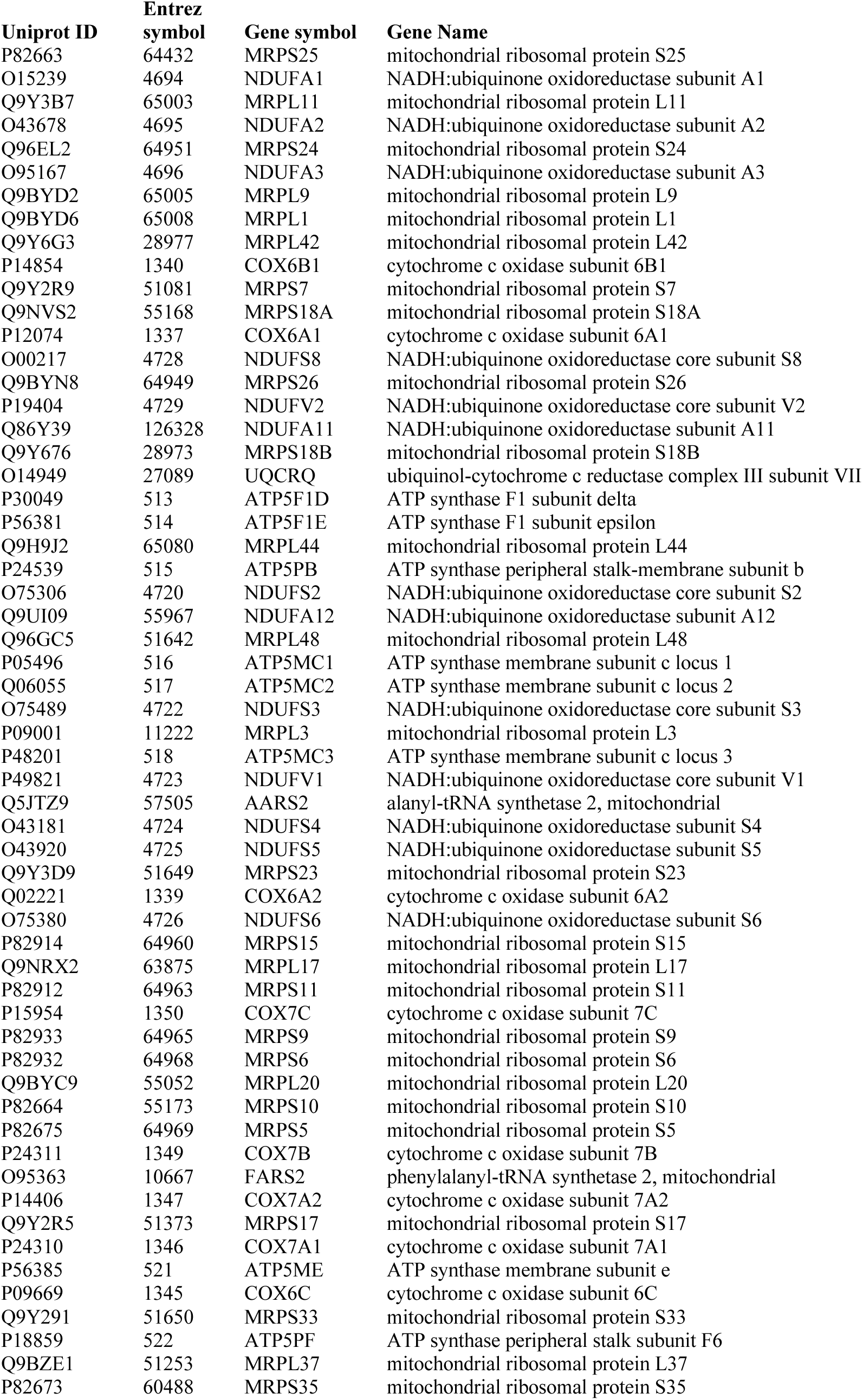
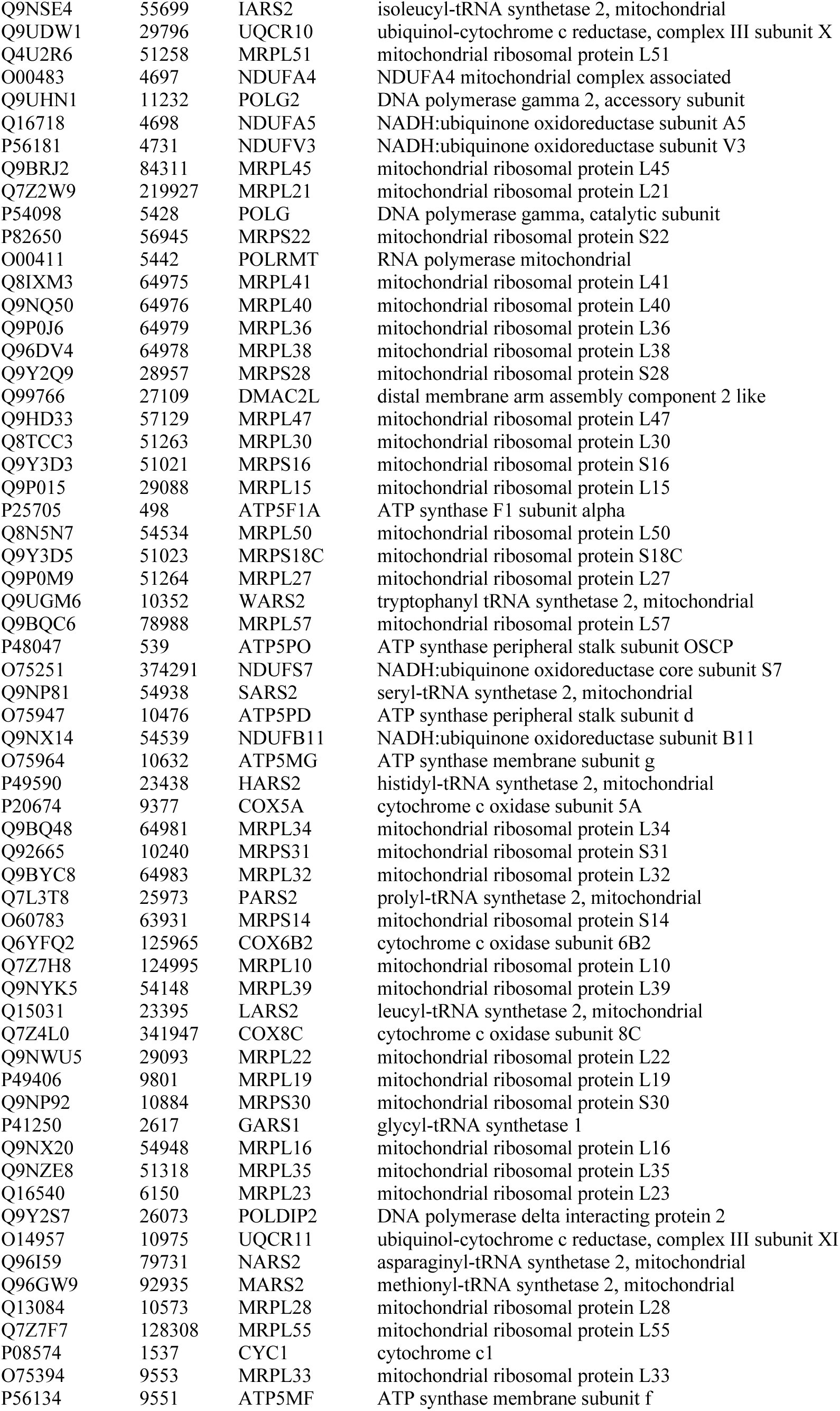
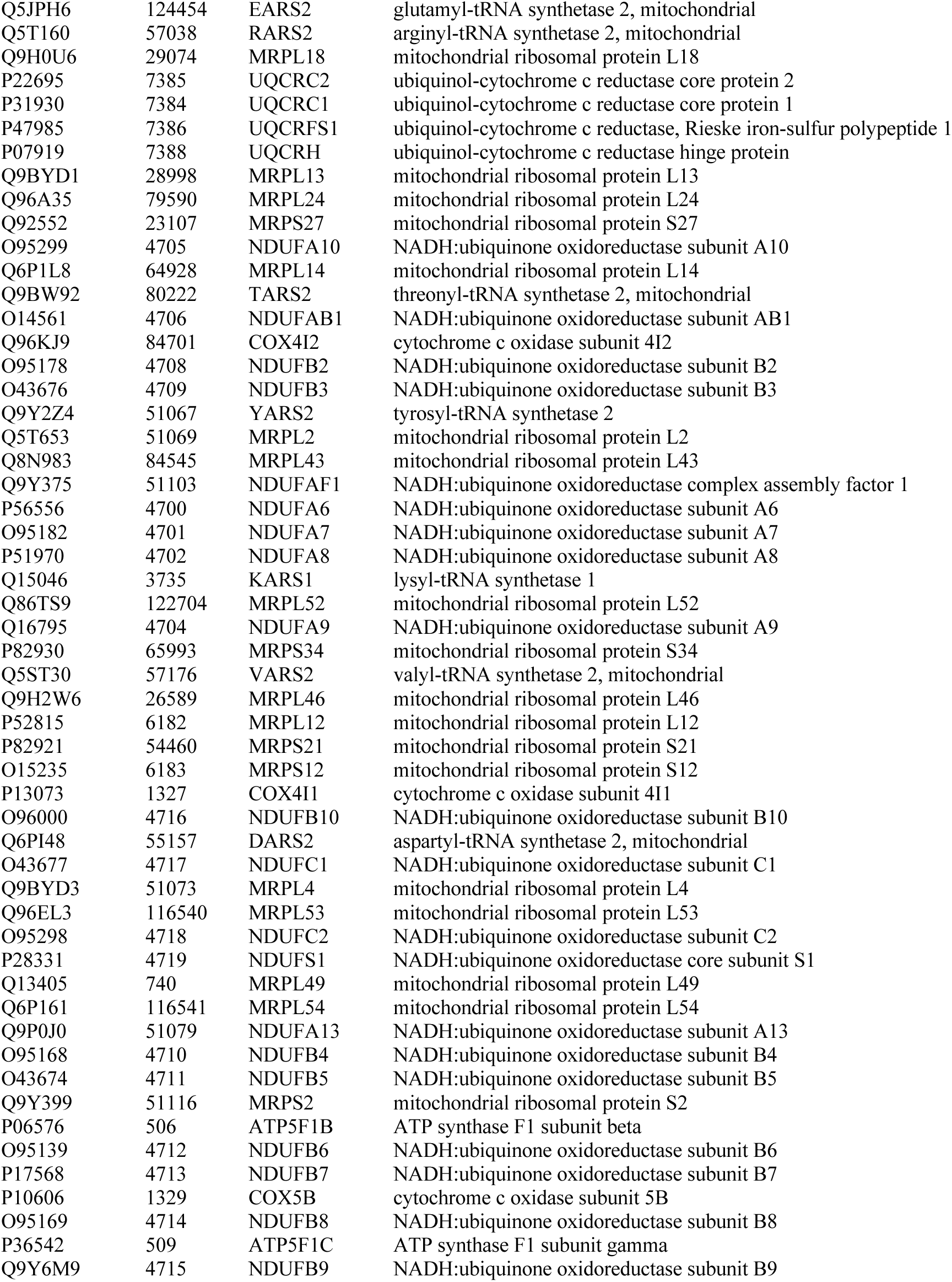
List of human interacting n-mt genes used in this study, modified from [10].

**Table S3.** Pre-filtered AIMs and genes (in parentheses) belonging to each gene class. The orthogroup row indicates how many orthogroups containing X. birchmanni protein sequences were found for each gene class.

| Population | Interacting n-mt | Non-interacting n-mt | Non-n-mt |
| --- | --- | --- | --- |
| Orthogroups | 159 | 822 | 15,545 |
| CALL AIMs (unique genes) | 954 (148) | 8,580 (883) | 287,834 (19,312) |
| CHAF AIMs (unique genes) | 952 (148) | 8,575 (883) | 287,592 (19,307) |
| HUEX-STAC AIMs (unique genes) | 997 (162) | 9,399 (947) | 319,679 (20,544) |

**Table S4.**
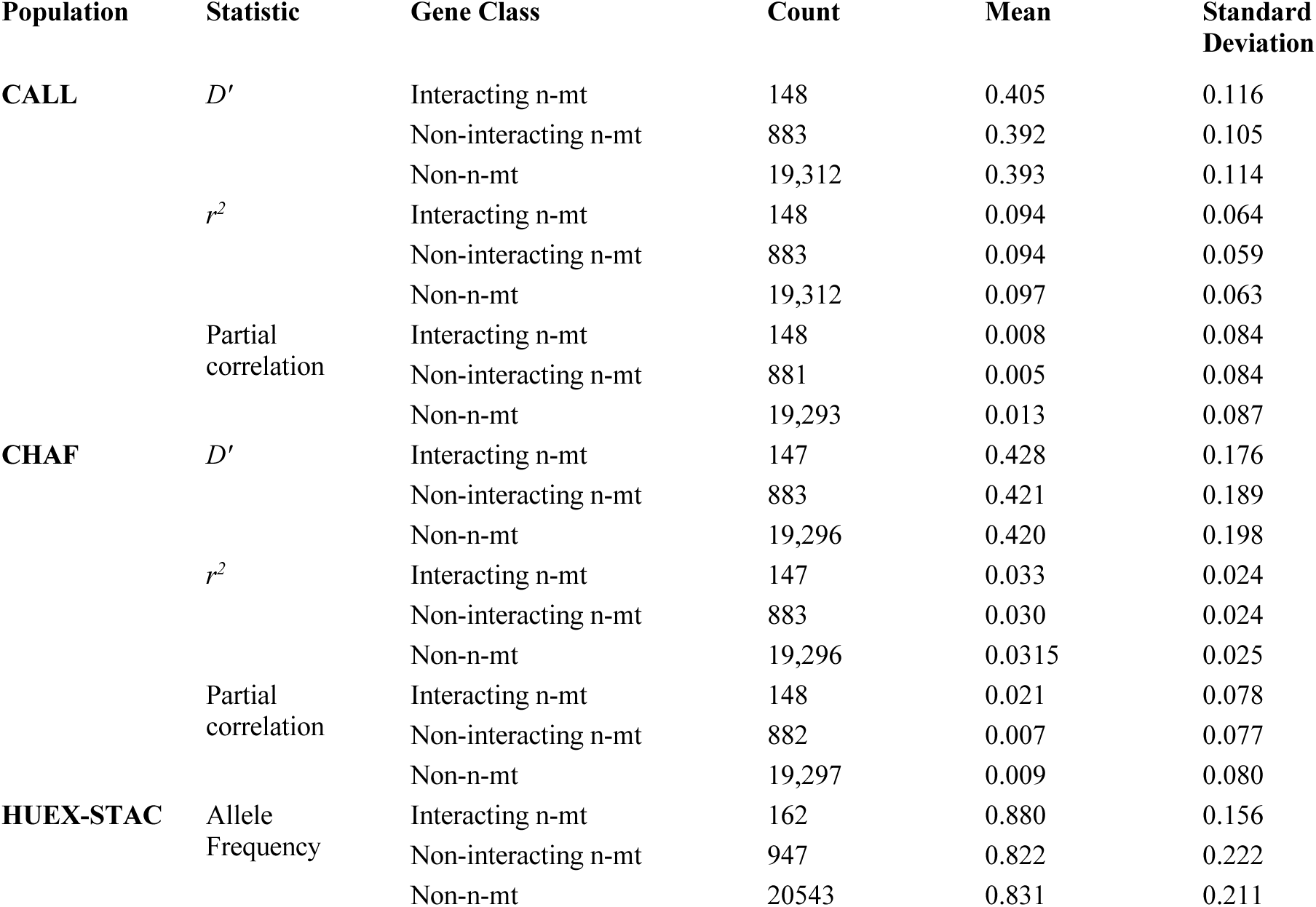
Summary statistics of mitonuclear association values for each hybrid population.

**Table S5.**
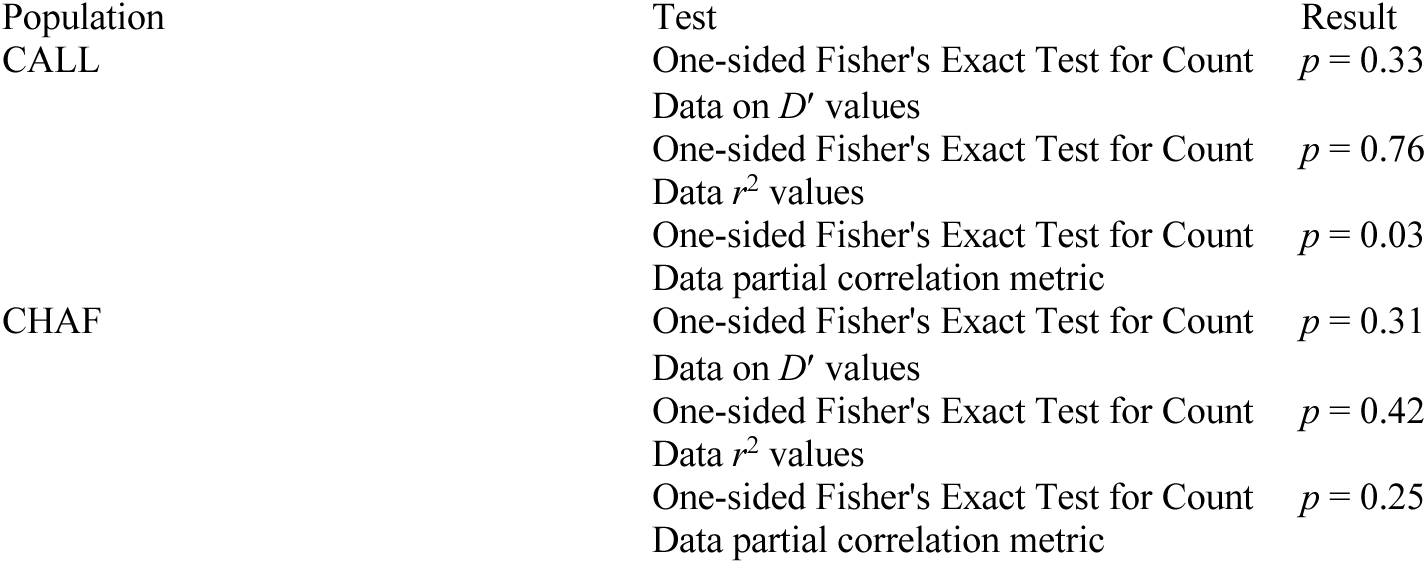
Results of Fisher’s exact test for enrichment of interacting n-mt genes in top 1% of genome-wide association statistics.

**Table S6.**
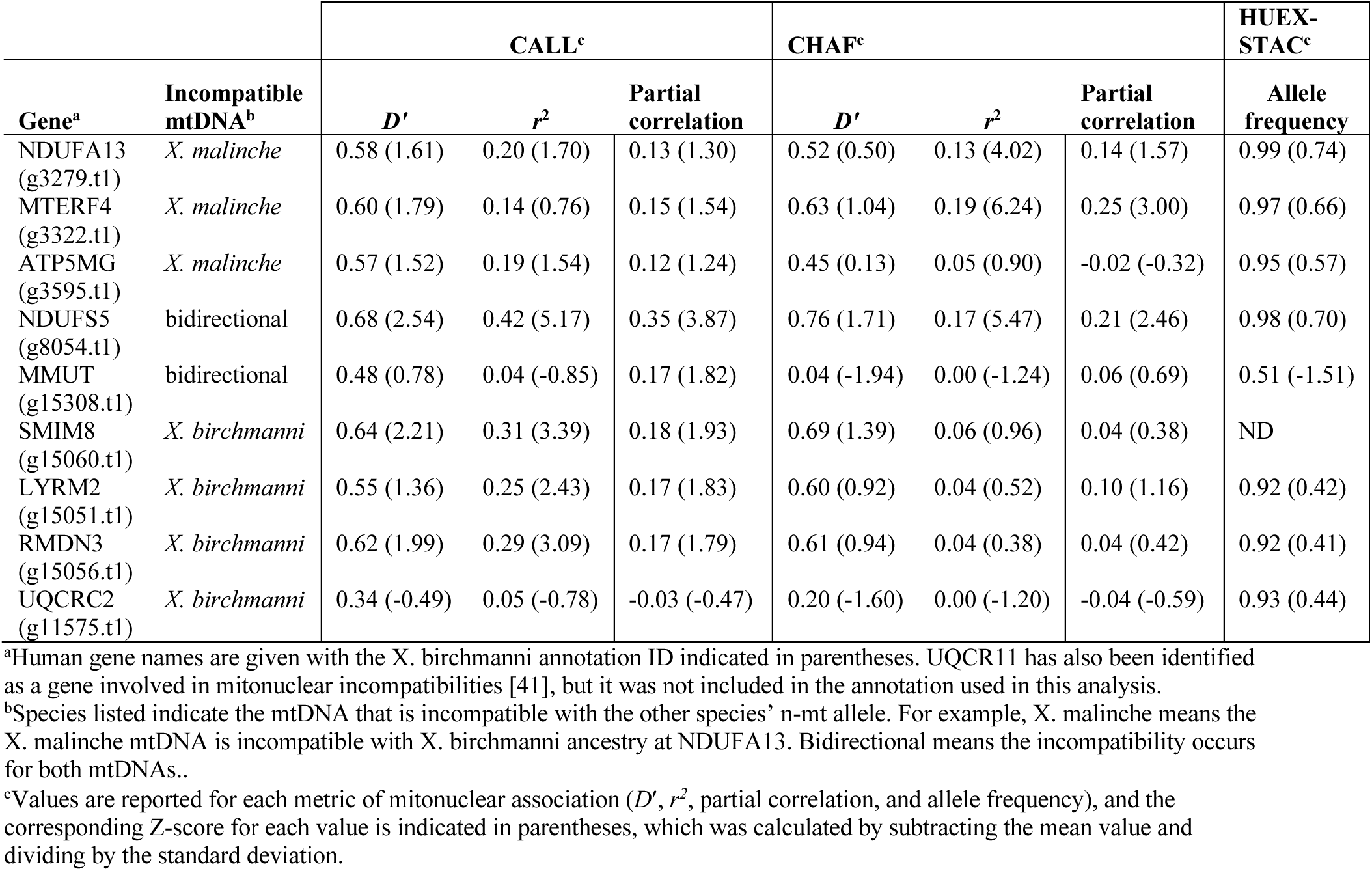
Measures of mitonuclear association for incompatible n-mt genes previously identified by Robles et al. [1].

**Table S7.** Summary statistics for the non-synonymous substitution model. NS is the count of non-synonymous substitutions per coding sequence divided by the coding sequence length.

| | Gene class | Number of genes | Mean NS ± standard deviation |
| --- | --- | --- | --- |
| CALL | Interacting n-mt | 63 | 0.00085 ± 0.0012 |
| | Non-interacting n-mt | 473 | 0.00051 ± 0.0007 |
| | Non-n-mt | 11330 | 0.00053 ± 0.0007 |
| CHAF | Interacting n-mt | 63 | 0.00085 ± 0.0012 |
| | Non-interacting n-mt | 473 | 0.00051 ± 0.0007 |
| | Non-n-mt | 11324 | 0.00053 ± 0.0007 |
| HUEX-STAC | Interacting n-mt | 60 | 0.00089 ± 0.0011 |
| | Non-interacting n-mt | 461 | 0.00046 ± 0.0007 |
| | Non-n-mt | 10587 | 0.00041 ± 0.0006 |

**Table S8.** Type III ANOVA results for non-synonymous substitution models.

|  | Predictor variables | p-value |
| --- | --- | --- |
| *D'* |  |  |
| CALL | **Intercept** | **< 2e-16** |
|  | Gene class | 0.40 |
|  | Non-synonymous substitutions / CDS (NS) | 0.16 |
|  | Gene class * NS | 0.11 |
| CHAF | **Intercept** | **< 2e-16** |
|  | Gene class | 0.93 |
|  | Non-synonymous substitutions / CDS (NS) | 0.54 |
|  | Gene class * NS | 0.35 |
| *r*<sup>2</sup> |  |  |
| CALL | **Intercept** | **< 2e-16** |
|  | **Gene class** | **0.042** |
|  | Non-synonymous substitutions / CDS (NS) | 0.42 |
|  | **Gene class \* NS** | **0.011** |
| CHAF | **Intercept** | **< 2e-16** |
|  | Gene class | 0.80 |
|  | Non-synonymous substitutions / CDS (NS) | 0.12 |
|  | Gene class * NS | 0.085 |
| Partial correlation |  |  |
| CALL | **Intercept** | **< 2e-16** |
|  | Gene class | 0.062 |
|  | Non-synonymous substitutions / CDS (NS) | 0.23 |
|  | Gene class * NS | 0.073 |
| CHAF | **Intercept** | **< 2e-16** |
|  | Gene class | 0.67 |
|  | Non-synonymous substitutions / CDS (NS) | **0.0028** |
|  | Gene class * NS | 0.67 |
| Allele Frequency |  |  |
| HUEX- | **Intercept** | **< 2e-16** |
| STAC | Gene class | 0.53 |
|  | **Non-synonymous substitutions / CDS (NS)** | **0.0034** |
|  | Gene class * NS | 0.83 |

Table S9. Results of summary() function testing the difference between interacting n-mt genes’ relationship between non-synonymous substitutions and mitonuclear LD and the non-n-mt genes’ relationship. Please see attached excel.

**Table S10.** Influential points of non-synonymous substitution analysis.

| Mitonuclear LD metric | Influential genes | Mitonuclear LD value | Normalized non-synonymous substitution count | Removed significance of interaction?<sup>a</sup> | Removed significance of interacting n-mt interaction?<sup>b</sup> |
| --- | --- | --- | --- | --- | --- |
| CALL |  |  |  |  |  |
| $r^2$ | *NDUFS5* | 0.42 | 0.0031 | Y | Y |
| Partial correlation | *NDUFS5* | 0.35 | 0.0031 | Y | Y |
| CHAF |  |  |  |  |  |
| $r^2$ | *NDUFS5* | 0.17 | 0.0031 | Y | Y |
|  | *NDUFA13* | 0.13 | 0.0023 | Y | Y |
|  | *ATP5MG* | 0.45 | 0.0064 | Y | N |
|  | *LOC102221627<sup>c</sup>* | 0.003 | 0.0031 | N | N |

The above points were identified as influential (i.e., significantly shaping the slope of the regression) in a model of mitonuclear LD vs an interaction between gene class and non-synonymous substitutions per coding sequence length.
<sup>a</sup>When this gene was removed from the model, did the interaction between gene class and normalized non-synonymous substitution count become non-significant?
<sup>b</sup>When this gene was removed from the model, did the interacting n-mt gene class no longer have a significantly sharper slope than the non-n-mt gene class?
<sup>c</sup>LOC102221627 is annotated as “cytochrome b-c1 complex subunit 8”

